# Critical roles for ‘housekeeping’ nucleases in Type III CRISPR-Cas immunity

**DOI:** 10.1101/2022.07.18.500432

**Authors:** Lucy Chou-Zheng, Asma Hatoum-Aslan

## Abstract

CRISPR-Cas systems are a family of adaptive immune systems that use small CRISPR RNAs (crRNAs) and CRISPR-associated (Cas) nucleases to protect prokaryotes from invading plasmids and viruses (i.e. phages). Type III systems launch a multi-layered immune response that relies upon both Cas and non-Cas cellular nucleases, and although the functions of Cas components have been well described, the identities and roles of non-Cas participants remain poorly understood. Previously, we showed that the Type III-A CRISPR-Cas system in *Staphylococcus epidermidis* employs two degradosome-associated nucleases, PNPase and RNase J2, to promote crRNA maturation and eliminate invading nucleic acids (Chou-Zheng and Hatoum-Aslan, 2019). Here, we identify RNase R as a third ‘housekeeping’ nuclease critical for immunity. We show that RNase R works in concert with PNPase to complete crRNA maturation, and identify specific interactions with Csm5, a member of the Type III effector complex, which facilitate nuclease recruitment/stimulation. Further, we demonstrate that RNase R and PNPase are required to maintain robust anti-plasmid immunity, particularly when targeted transcripts are sparse. Altogether, our findings expand the known repertoire of accessory nucleases required for Type III immunity and highlight the remarkable capacity of these systems to interface with diverse cellular pathways to ensure successful defense.

## Introduction

CRISPR-Cas (<u>C</u>lustered regularly-interspaced <u>s</u>hort <u>p</u>alindromic repeats-<u>C</u>RISPR <u>a</u>ssociated) systems are adaptive immune systems in prokaryotes that use small CRISPR RNAs (crRNAs) in complex with Cas nucleases to sense and degrade foreign nucleic acids (Barrangou et al. 2007; Jansen et al. 2002; Hille et al. 2018). The CRISPR-Cas pathway generally occurs in three stages—adaptation, crRNA biogenesis, and interference. During adaptation, Cas nucleases clip out short sequences (known as ‘protospacers’) from invading nucleic acids and integrate them into the CRISPR locus as ‘spacers’ in between short DNA repeats. During crRNA biogenesis, the repeat-spacer array is transcribed as a long precursor crRNA (pre-crRNA) which is subsequently processed within repeats to generate mature crRNAs that each bear a single spacer sequence. Mature crRNAs combine with one or more Cas nucleases to form effector complexes which, during interference, detect and cleave matching nucleic acid invaders. Although all CRISPR-Cas systems follow this general pathway, they exhibit striking diversity in the composition of their effector complexes and mechanisms of action. Accordingly, they have been divided into two classes, six Types (I-VI), and over 30 subtypes (Makarova, Wolf, et al. 2020; Koonin and Makarova 2022).

Type III CRISPR-Cas systems are the most closely related to the ancestral system from which all Class I systems have evolved and are arguably the most complex (Mohanraju et al. 2016; Koonin and Makarova 2022). Type III systems typically utilize multi-subunit effector complexes which recognize foreign RNA and coordinate a sophisticated immune response that results in the destruction of the invading RNA and DNA. Of the six subtypes currently identified (A-F), Types III-A and III-B are the best characterized. In these systems, crRNA binding to a complementary transcript triggers at least three catalytic activities by members of the effector complex: target RNA shredding by Cas7/Csm3/Cmr4 (Hale et al. 2009; Staals et al. 2013; 2014; Samai et al. 2015; Tamulaitis et al. 2014), nonspecific DNA degradation by Cas10 (Samai et al. 2015; Kazlauskiene et al. 2016; Estrella, Kuo, and Bailey 2016; Liu, Iavarone, and Doudna 2017; Elmore et al. 2016), and Cas10-catalyzed production of cyclic-oligoadenylates (cOAs), second-messenger molecules which bind and stimulate accessory nucleases outside of the effector complex (Niewoehner et al. 2017; Kazlauskiene et al. 2017; Han et al. 2018; Nasef et al. 2019). Such accessory nucleases typically possess CRISPR-associated Rossman Fold (CARF) domains to which cOAs bind and are encoded within or proximal to the Type III CRISPR-Cas locus (Shmakov et al. 2018; Shah et al. 2019; Makarova, Timinskas, et al. 2020). Indeed, recent studies have relied upon these two features to discover new cOA-responsive accessory nucleases and validate their contributions to Type III defense (Han et al. 2018; Athukoralage et al. 2019; McMahon et al. 2020; Rostøl et al. 2021; Zhu et al. 2021). However, as the list of CRISPR-associated accessory nucleases continues to grow, the identities and contributions of non-Cas participants in Type III immunity remain poorly understood.

Our previous work showed that the Type III-A CRISPR-Cas system in *Staphylococcus epidermidis* (herein referred to as CRISPR-Cas10) employs the ‘housekeeping’ nucleases PNPase and RNase J2 during multiple steps in the immunity pathway (Walker et al. 2017; Chou-Zheng and Hatoum-Aslan 2019) (Fig. 1 A-C). During crRNA biogenesis, Cas6 cleaves pre-crRNAs within repeats to generate 71 nucleotide (nt) intermediates (Hatoum-Aslan et al. 2014), and these intermediates are processed on their 3’-ends by PNPase and one or more unidentified nuclease(s) to produce mature species of 43, 37, and 31 nt in length (Chou-Zheng and Hatoum-Aslan 2019). We also showed that PNPase helps to prevent phage nucleic acid accumulation during an active infection, implying a direct role for PNPase during interference. While searching for additional maturation nuclease(s), we fortuitously identified RNase J2 as another player in the pathway; however, while it has little/no effect on crRNA maturation, RNase J2 is essential for interference against phage and plasmid invaders. Notably, PNPase and RNase J2 are members of the RNA degradosome, a highly-conserved complex of ribonucleases, helicases, and metabolic enzymes primarily involved in RNA processing and decay (Tejada-Arranz, de Crecy-Lagard, and de Reuse 2020). Our original observation that these nucleases co-purify in trace amounts with the Cas10-Csm complex in *S. epidermidis* (Walker et al. 2017) led to the discovery of their additional contributions to CRISPR-Cas defense.

**Figure 1.**
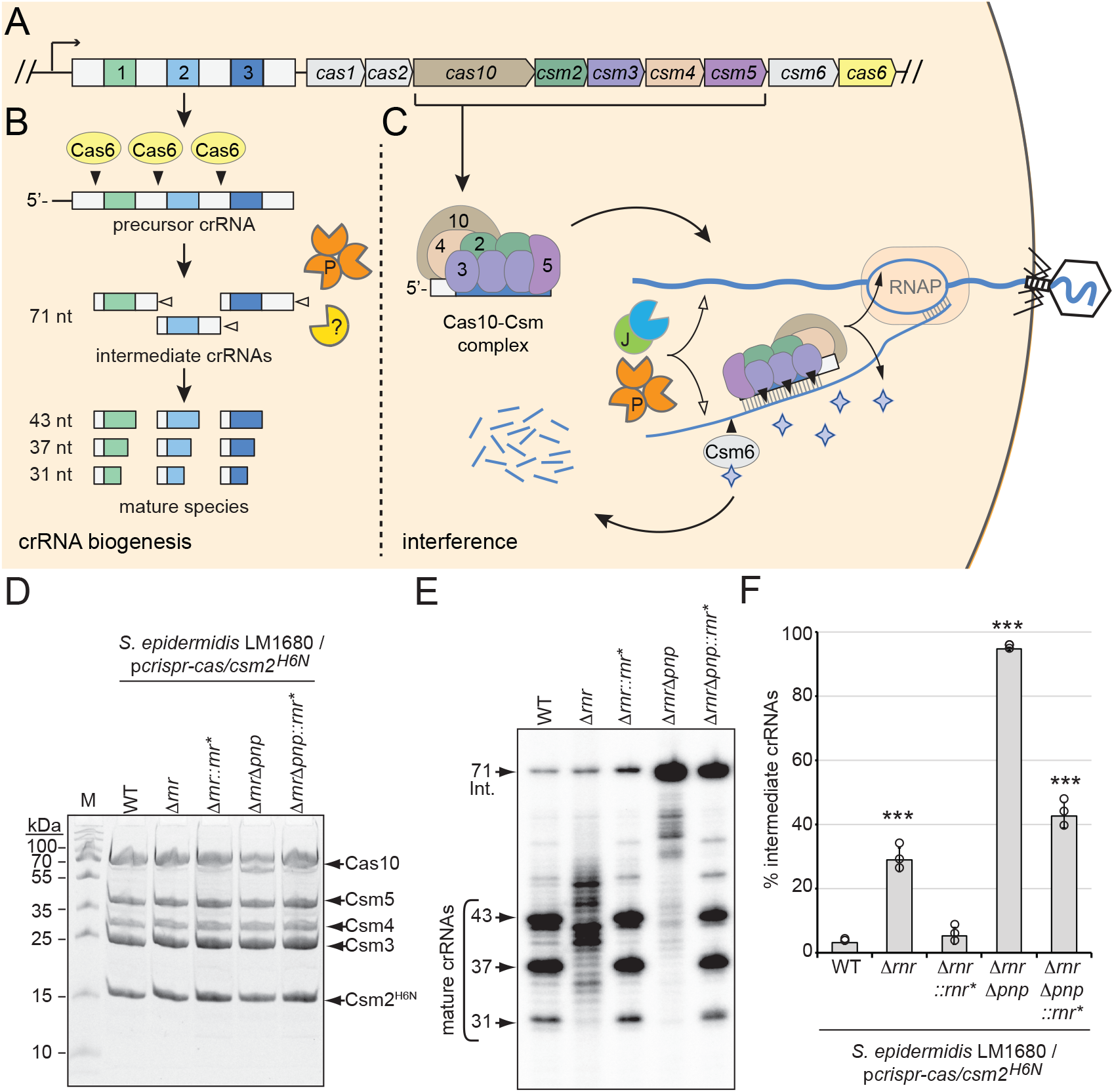
RNase R and PNPase are necessary for crRNA maturation in the cell (A) The type III-A CRISPR-Cas system (herein referred to as CRISPR-Cas10) in *S. epidermidis* RP62a encodes three spacers (colored squares), four repeats (light grey squares) and nine CRISPR-associated (*cas* and *csm*) genes (colored pentagons). (B) During crRNA biogenesis, the repeat-spacer array is transcribed into a precursor crRNA and processed into mature species in two steps. In the first step, the endoribonuclease Cas6 cleaves within repeat sequences to generate intermediate crRNAs of 71 nt in length. In the second step, intermediates are trimmed on their 3’-ends by PNPase and other unknown nuclease(s) which are the subject of this study. These activities generate mature crRNAs that range from 43 to 31 nt in length. (C) Mature crRNAs associate with Cas10, Csm2, Csm3, Csm4, and Csm5 in various stoichiometries to form the Cas10-Csm effector complex. Interference is initiated when the effector complex binds to invading transcripts that bear complementarity to the crRNA. During interference, invading DNA and RNA are degraded by CRISPR-associated (Cas) and non-Cas nucleases (see text for details). Filled triangles illustrate events catalyzed by Cas enzymes, and open triangles illustrate events catalyzed by non-Cas nucleases. P, PNPase; J, RNase J1/J2; RNAP, RNA polymerase. Purple stars represent cyclic oligoadenylate molecules produced by Cas10. (D) Cas10-Csm complexes extracted from indicated *S. epidermidis* LM1680 strains bearing p*crispr-cas/csm2*^*H6N*^ are shown. The plasmid p*crispr-cas* encompasses the entire CRISPR-Cas10 system with a 6-His tag on the N-terminus of Csm2. Whole cell lysates from indicated strains were subjected to Ni^2+^ affinity chromatography, and purified complexes were resolved in an SDS-PAGE gel and visualized with Coomassie G-250 staining. M, denaturing protein marker; kDa, kilodalton. See Figure 1-source data 1. (E) Total crRNAs associated with Cas10-Csm complexes in panel D are shown. Complex-bound crRNAs were extracted from complexes, radiolabeled at their 5’-ends, and resolved on a denaturing gel. See Figure 1-source data 2. (F) Fractions of complex-bound intermediate crRNAs relative to total crRNAs are shown for indicated strains. The percent intermediate crRNAs represents the ratio of the intermediate (71 nt) band density to the sum of band densities of the major crRNA species (71, 43, 37, and 31 nt). Data shown represents an average of 3 independent trials (±S.D). A two-tailed t-test was performed to determine significance and *** indicates *p* < 0.0005. See Figure 1-source data 3.

Here, we sought to complete the crRNA maturation pathway in *S. epidermidis* and discovered that RNase R is the second (and final) nuclease necessary for the process. We demonstrate that RNase R works in concert with PNPase to catalyze crRNA maturation in a purified system, and these enzymes work synergistically in the cell to maintain robust anti-plasmid immunity. Further, we identified specific interactions between these ‘housekeeping’ nucleases and Csm5 (a member of the Cas10-Csm complex within the Cas7 group) which facilitate their recruitment and/or stimulation. Altogether, our findings expand the known repertoire of non-Cas nucleases that facilitate Type III CRISPR-Cas defense and highlight the remarkable capacity of this system to interface with diverse non-defense cellular pathways to maintain robust immunity.

## Results

### RNase R and PNPase are necessary for crRNA maturation in the cell

Previously, we showed that an in-frame deletion of *pnp* (which encodes PNPase) in *S. epidermidis* causes loss of about half of the mature crRNA species and significant (∼10-fold) accumulation of intermediates (Chou-Zheng and Hatoum-Aslan 2019), indicating that one or more additional nucleases contribute to crRNA maturation. We also showed previously that PNPase co-purifies with the Cas10-Csm complex in sub-stoichiometric amounts along with at least five additional cellular nucleases which serve as maturation nuclease candidates—RNase J1, RNase J2, Cbf1, RNase R, and RNase III (Walker et al. 2017). Given that crRNA maturation relies upon 3’-5’ exonuclease activity, here we sought investigate the two remaining nucleases in the list that possess this function—Cbf1 and RNase R. Unfortunately, repeated attempts to delete *cbf1* from *S. epidermidis* failed, suggesting it may be essential for cell viability under standard laboratory growth conditions. However, *rnr* (which encodes RNase R) was readily deleted in the clinical isolate *S. epidermidis* RP62a (Christensen, Baddour, and Simpson 1987) as well as in *S. epidermidis* LM1680, a mutant variant of RP62a that has lost the CRISPR-Cas system (Jiang et al. 2013) (Figure 1-figure supplement 1 A and B). To determine the extent to which RNase R contributes to crRNA maturation *in vivo*, a plasmid that encodes the Type III-A CRISPR-Cas system of RP62a, p*crispr-cas/csm2*^*H6N*^ (Hatoum-Aslan et al. 2013), was introduced into *S. epidermidis* LM1680/Δ*rnr*. Importantly, this construct encodes a 6-Histidine (6-His) tag on the N-terminus of Csm2 which allows for complex purification via Ni^2+^-affinity chromatography. Cas10-Csm complexes were subsequently purified from the wild-type (WT) and Δ*rnr* strains (Fig. 1D), crRNAs were further purified from the complexes and visualized (Fig. 1E), and fractions of intermediate species were quantified (Fig. 1F). This experiment revealed that deletion of RNase R alone causes complete loss of precisely-processed mature species and production of crRNAs with a range of aberrant lengths, as well as moderate accumulation of 71 nt intermediates. To confirm that the loss of RNase R is responsible for this phenotype, we returned *rnr* to its native locus in the genome to generate *S. epidermidis* LM1680/Δ*rnr*::*rnr\**—in this strain, silent mutations were introduced into *rnr* to distinguish the knock-in strain from original WT (Figure 1-figure supplement 1 C and D). The same assays were then repeated and showed that crRNAs in the complemented strain have sizes similar to those found in WT (Fig. 1 D-F). These results demonstrate that RNase R is necessary for crRNA maturation *in vivo*.

In order to determine the extent to which other nuclease(s) may contribute to crRNA maturation in the absence of RNase R and PNPase, we created and tested a double-knockout. Specifically, *rnr* was deleted from the LM1680/Δ*pnp* strain (Chou-Zheng and Hatoum-Aslan 2019) to generate LM1680/Δ*rnr*Δ*pnp*, and crRNAs within the complexes were examined. The results showed a complete loss of mature species in the double mutant, with ∼95% of crRNAs trapped in the intermediate state (Fig. 1 E and F). In addition, when *rnr* is returned to the double mutant (i.e. in LM1680/Δ*rnr*Δ*pnp::rnr\**), some mature crRNAs are recovered with significant accumulation of the 71 nt intermediates, similar to the phenotype observed in LM1680/Δ*pnp* (Chou-Zheng and Hatoum-Aslan 2019). Altogether, these data demonstrate that RNase R and PNPase are likely the primary drivers of crRNA maturation in the cell.

### RNase R and PNPase are sufficient to catalyze crRNA maturation in a purified system

Given that deletion of RNase R on its own causes complete loss of precisely-processed mature species *in vivo*, while deletion of PNPase still allows for some maturation to occur, the possibility exists that RNase R alone might be sufficient to catalyze maturation to completion in a purified system (i.e. in the absence of nonspecific cellular RNA substrates). Conversely, it is also possible that other nucleases in the cell (such as Cbf1) might contribute to crRNA maturation. Indeed, Cbf1 (also called YhaM) is known to work together with other 3’-5’ exonucleases in the cell to help clear RNA decay intermediates (Broglia et al. 2020). To determine the extent to which these housekeeping nucleases catalyze crRNA maturation on their own, we performed nuclease assays with purified components (Fig. 2 A). In these assays, Cas10-Csm complexes loaded with 71 nt intermediate crRNAs (Cas10-Csm (71)) were purified from LM1680/Δ*rnr*Δ*pnp* and combined with purified RNase R, PNPase, and/or Cbf1 (Fig. 2 B). After 30 minutes of incubation with appropriate divalent metals, crRNAs were extracted from the complexes and visualized (Fig. 2 C and D). As expected, the Cas10-Csm (71) complex is unable to catalyze crRNA maturation on its own. Interestingly, Cbf1 is also incapable of cleaving intermediate crRNAs associated with the complex. In contrast, RNase R and PNPase each cause partial processing of crRNA intermediates on their 3’-ends, with cleavage patterns similar to those observed when one or the other nuclease is deleted from cells—addition of PNPase produces a pattern of crRNA lengths similar to that seen in LM1680/Δ*rnr* cells, and addition of RNase R into the reaction generates crRNA lengths similar to those extracted from LM1680/Δ*pnp* cells (compare Figs. 1E and 2C). Even when given up to 60 minutes in the *in vitro* assay, RNase R alone is unable to process about half of the intermediate crRNAs (Figure 2-figure supplement 1). However, when RNase R and PNPase are combined in the reaction, the majority of crRNA intermediates are processed to the appropriate mature lengths (43, 37, and 31 nt) with proportions bearing a striking resemblance to those observed when crRNAs are purified from WT cells (Fig. 2 C and D). Taken together, our data demonstrate that RNase R and PNPase are both necessary and sufficient to process intermediate crRNAs associated with the Cas10-Csm complex to achieve their final mature lengths.

**Figure 2.**
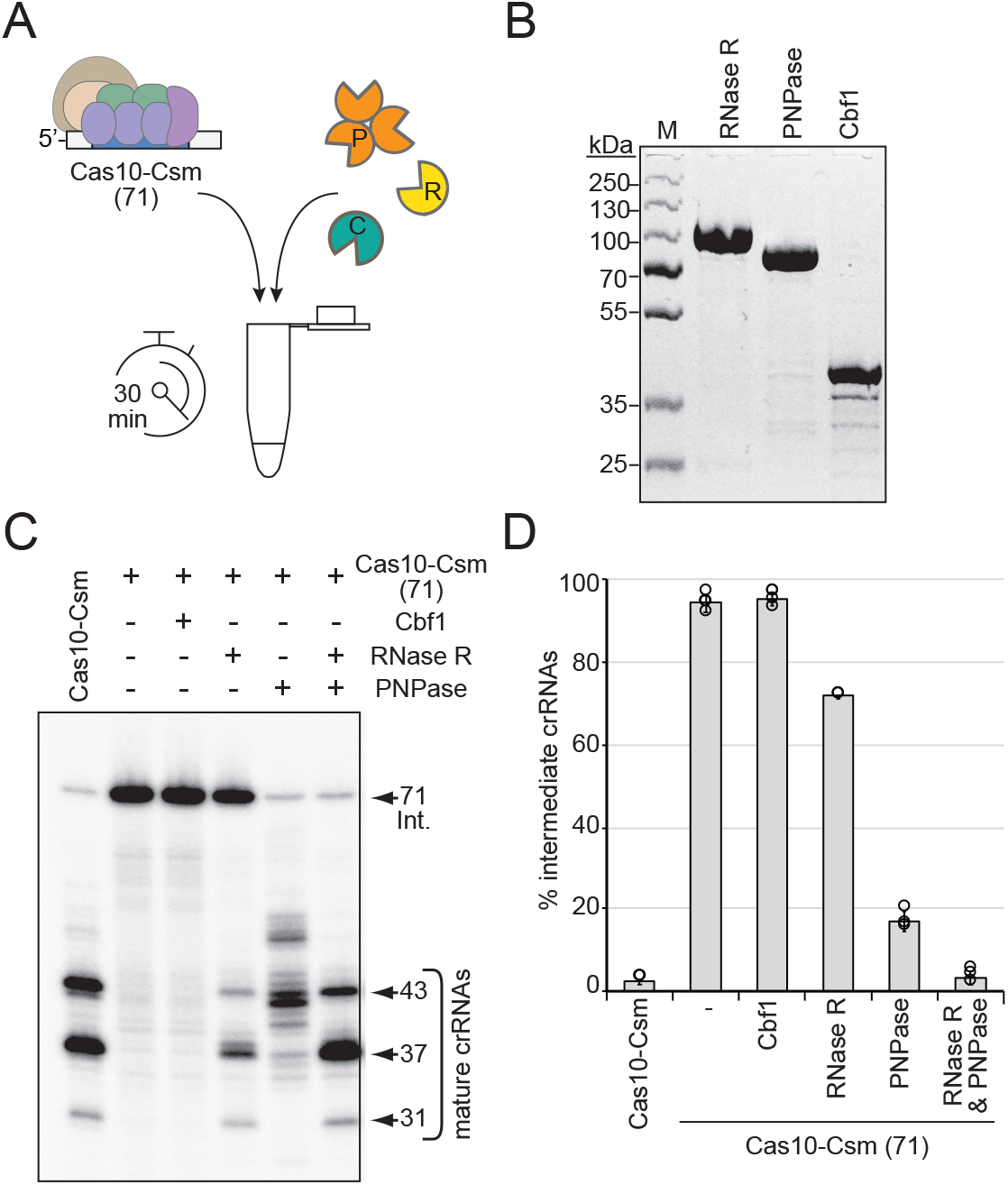
RNase R and PNPase are sufficient to complete crRNA maturation in A B a purified system. (A) Illustration of experimental flow of the crRNA maturation nuclease assay. P, PNPase; R, RNase R; C, Cbf1; Cas10-Csm (71), Cas10-Csm complexes purified from *S. epidermidis* LM1680Δ*pnp*Δ*rnr*. (B) Purified recombinant exonucleases RNase R, PNPase, and Cbf1 are shown. Proteins were resolved in an SDS-PAGE gel and visualized with Coomassie G-250 staining. M, denaturing protein marker. kDa, kilodalton. See Figure 2-source data 1. (C) Cas10-Csm (71) complexes were incubated with indicated nucleases for 30 minutes at 37°C. After digestion, crRNAs were extracted from the complexes, radiolabeled at their 5’-ends, and resolved on a denaturing gel. The leftmost lane shows crRNAs extracted from Cas10-Csm complexes purified from WT cells as a control. See also Figure 2-figure supplement 1 and Figure 2-source data 2. (D) Quantification of complex-bound intermediate crRNAs (relative to total crRNAs) following crRNA maturation assays. The data represent an average of 2–4 independent trials (±S.D). See Figure 2-source data 3.

### Csm5 interacts with RNase R

We next considered the mechanism of RNase R recruitment to the Cas10-Csm complex. Previously, we showed that Csm5 (a member of the complex within the Cas7 group) directly interacts with PNPase in a purified system (Walker et al. 2017), and since deletion of *csm5* causes complete loss of crRNA maturation while allowing for the remainder of the complex to form (Hatoum-Aslan et al. 2013), we reasoned that RNase R recruitment is also likely to be facilitated by Csm5. To test this, RNase R and Csm5 were resolved alone and combined in a native polyacrylamide gel, which separates proteins on the basis of size and charge. As expected, Csm5 fails to enter into the gel at near-neutral pH due to its basic isoelectric point (pI, Table S1) and resulting positive charge in the native running conditions, while RNase R migrates into the native gel and shows up as a band following Coomassie staining (Fig. 3A and B). Consistent with previous observations, RNase R appears to self-associate *in vitro* (Zuo and Deutscher 2001), which accounts for its shallow migration into the gel. Nonetheless, we observed that the addition of increasing amounts of Csm5 to RNase R causes the band to shift upward, indicating that the two proteins interact. This interaction is likely weak/transient because Csm5 must be added in excess (up to 9:1) to observe a noticeable band shift. To confirm that the interaction is specific to RNase R, we repeated the same assay using bovine serum albumin (BSA), which also has an acidic isoelectric point (Table S1), and no such shift was observed (Figure 3-figure supplement 1). These data suggest that Csm5 facilitates recruitment of RNase R to the Cas10-Csm complex.

**Figure 3.**
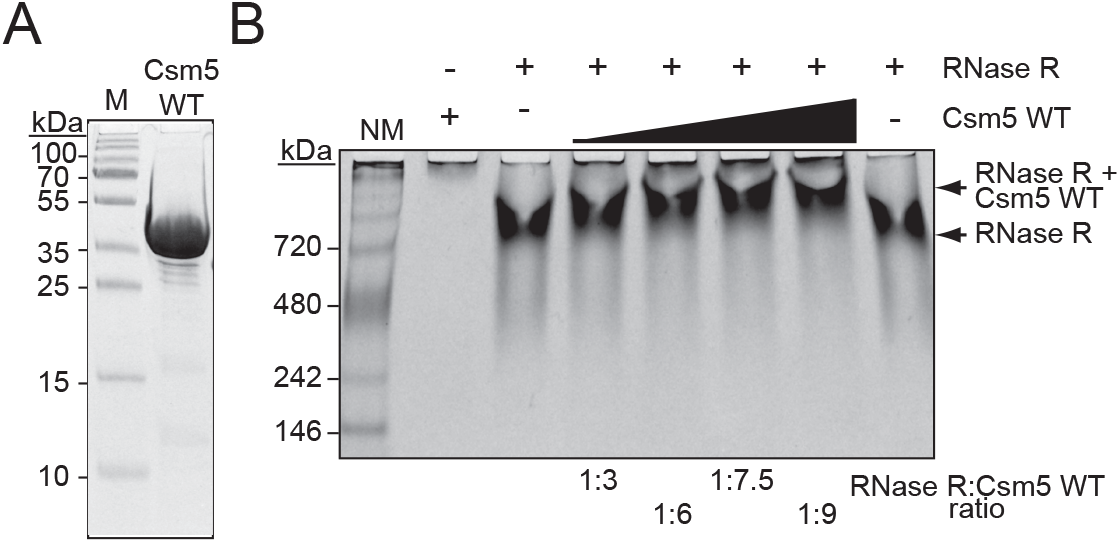
Csm5 interacts with RNase R. (A) Purified recombinant WT Csm5 is shown. The protein was resolved in an SDS-PAGE gel and visualized using Coomassie G-250 staining. M, denaturing protein marker; kDa, kilodalton. See Figure 3-source data 1. (B) Native gel showing RNase R resolved with increasing proportions of Csm5 WT. Shown is a representative of three independent trials. NM, native protein marker. See also Figure 3-figure supplement 1 and Figure 3-source data 2.

### Csm5 binds and stimulates PNPase through a predicted disordered region

Csm5 is about half the size of RNase R and PNPase (Table S1), and considering that PNPase functions as a trimer (Symmons, Jones, and Luisi 2000), we wondered how Csm5 provides binding sites for both proteins. One possibility is that the nuclease docking site(s) might be spread over multiple subunits of the Cas10-Csm complex, with Csm5 contributing to the bulk of the interaction(s). Another non-exclusive possibility is that both nucleases may be recruited by the same/overlapping binding site(s) on Csm5, with one or the other allowed to occupy the site at any given time. Such transient and dynamic interactions are known to occur with proteins bearing intrinsically disordered regions (IDRs), flexible polypeptides enriched with charged residues that have the capacity to bind multiple partners (Dyson and Wright 2005; Chakrabarti and Chakravarty 2022; Bigman, Iwahara, and Levy 2022). Indeed, Csm5 is enriched with positively-charged amino acids which confer its basic pI (Fig. 4A and Table S1). Based on these observations, we hypothesized that Csm5 mediates binding of RNase R and/or PNPase via one or more IDR(s).

**Figure 4.**
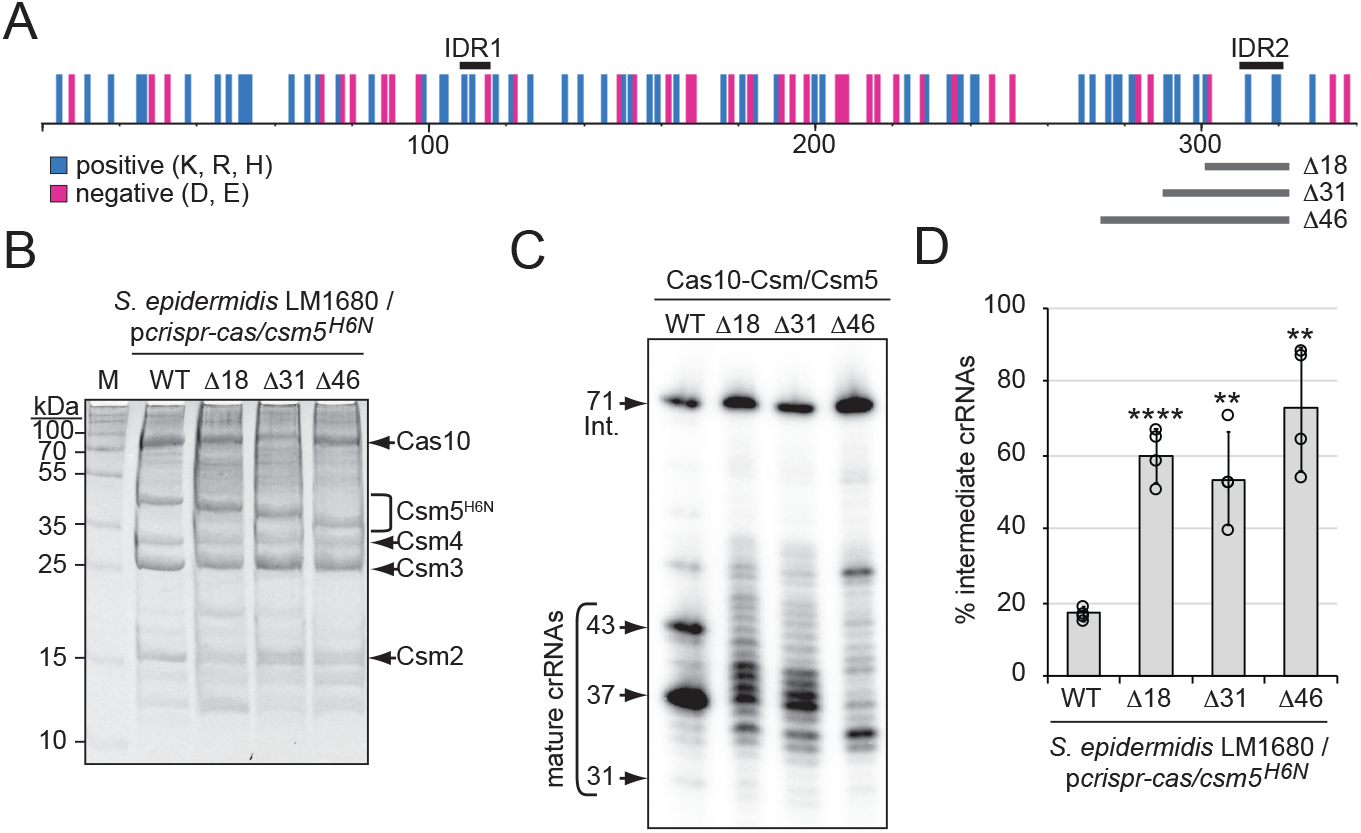
A predicted disordered region in Csm5 promotes crRNA maturation (A) Illustration showing the distribution of charged residues, predicted disordered regions, and truncations introduced in Csm5. Positions of charged residues (positive, cyan; negative, magenta) are shown as vertical bars. Predicted intrinsically disordered regions (IDR1 and IDR2) and regions that were truncated are delimited by black and grey horizontal bars above and below, respectively. K, lysine; R, arginine; H, histidine; D, aspartate; E, glutamate. See also Figure 4-figure supplement 1. (B) Cas10-Csm complexes with various Csm5 truncations are shown. Complexes were extracted from *S. epidermidis* LM1680 cells harboring p*crispr-cas*/*csm5*^*H6N*^, which has a 6-His tag on the N-terminus of Csm5 to confirm full complex assembly. Complexes were purified using Ni^2+^ affinity chromatography, resolved on and SDS-PAGE gel, and visualized with Coomassie G-250 staining. M, denaturing protein marker; kDa, kilodalton. See also Figure 4-source data 1. (C) Total crRNAs bound to indicated Cas10-Csm complexes were extracted, radiolabeled at their 5’-ends, and resolved on a denaturing gel. See also Figure 4-source data 2. (D) Fractions of complex-bound intermediate crRNAs relative to total crRNAs are shown for Csm5 truncation mutants. The percent of intermediate crRNAs represents the ratio of the intermediate (71 nt) band density to the sum of band densities of the major crRNA species (71, 43, 37, and 31 nt). The data represents an average of 4 independent trials (±S.D). A two-tailed t-test was performed to determine significance and *p-*values obtained were < 0.005 (**) and 0.00005 (****). See also Figure 4-source data 3.

To begin to test this hypothesis, we searched for predicted disordered regions in *S. epidermidis* Csm5 using the web-based tool PONDR (predictor of natural disordered regions), which uses neural networks to discriminate between ordered and disordered residues in a given protein (Romero, Obradovic, and Dunker 2004). This analysis revealed the presence of two putative disordered regions spanning residues 109-116 and 310-320 (here onward IDR1 and IDR2, respectively), with the latter having the higher probability for being disordered (Figure 4-figure supplement 1 A and B). These regions were next examined in relation to the distribution of charged residues across Csm5, and we noted that both IDRs encompass multiple positively-charged residues (Fig. 4A). To gain a better understanding of the structural context of the predicted IDRs, we examined the homologous residues in the experimentally determined structure of the Cas10-Csm complex from *Streptococcus thermophilus* (StCas10-Csm) (You et al. 2019). Homologous residues in Csm5 were identified in a Clustal pairwise sequence alignment and mapped back to the StCsm5 structure. We found that the StCsm5 subunit in the unbound complex (PDB ID 6IFN) has half of its amino acids within loop/coil structures (Figure 4-figure supplement 1 C, magenta), and the residues homologous to those comprising IDR2 in *S. epidermidis* Csm5 align well with a long loop structure in StCsm5 (Figure 4-figure supplement 1C, cyan). These observations lend support to the notion that Csm5 may possess one or more IDRs which play role(s) in nuclease recruitment.

To further test this hypothesis, we deleted the regions encoding IDR1 and IDR2 from *csm5* in a plasmid bearing the entire CRISPR-Cas system of *S. epidermidis* RP62a (p*crispr-cas/csm5*^*H6N*^) (Hatoum-Aslan et al. 2013). Importantly, the 6-His tag in this construct is located on Csm5 to allow us to rule out mutations that impact Csm5 stability and/or complex assembly. The plasmids were introduced into *S. epidermidis* LM1680 and cell lysates were subjected to Ni^2+^-affinity chromatography. We were unable to purify complexes when the IDR1 region in Csm5 was deleted (not shown), indicating that the residues comprising IDR1 might be important for Csm5 stability and/or complex assembly. In contrast, full complexes were recovered in the presence of three deletions spanning 18, 31, or 46 amino acids encompassing IDR2 (Fig. 4 A and B). Interestingly, crRNAs bound to these complexes exhibited a range of aberrant lengths with significant (>50%) accumulation of 71 nt intermediates (Fig. 4 C and D), suggesting that IDR2 within Csm5 may facilitate interactions with RNase R and/or PNPase.

We further tested for direct interactions using gel shift assays, in which Csm5Δ46 was purified (Fig. 5A), combined with RNase R or PNPase in different proportions, and resolved on native polyacrylamide gels. The results showed that while Csm5Δ46 maintains its interaction with RNase R (Figure 5-figure supplement 1), the mutant has little/no interaction with PNPase (Fig. 5B), indicating that the IDR2 region is essential for PNPase binding *in vitro*. Previously, we showed that in addition to the physical interaction between Csm5 and PNPase, there is also a functional interaction in which Csm5 stimulates PNPase’s nucleolytic activity (Walker et al. 2017). To determine the impact of IDR2 on PNPase activity, nuclease assays were performed in which a 31 nt ssRNA substrate was incubated with PNPase and/or Csm5 for increasing amounts of time. Consistent with previous results, we found that Csm5 alone has no impact on substrate length, PNPase (which is a dual polymerase-nuclease) causes both RNA extension and degradation, and when the two proteins are combined, a striking stimulation of PNPase’s nucleolytic activity occurs (Fig. 5C). Importantly, Csm5Δ46 fails to cause such stimulation, consistent with the loss of physical interaction between Csm5Δ46 and PNPase. Taken together, these results demonstrate that the IDR2 region of Csm5 plays a critical role in the recruitment and stimulation of PNPase, while the binding site for RNase R likely resides elsewhere in Csm5.

**Figure 5.**
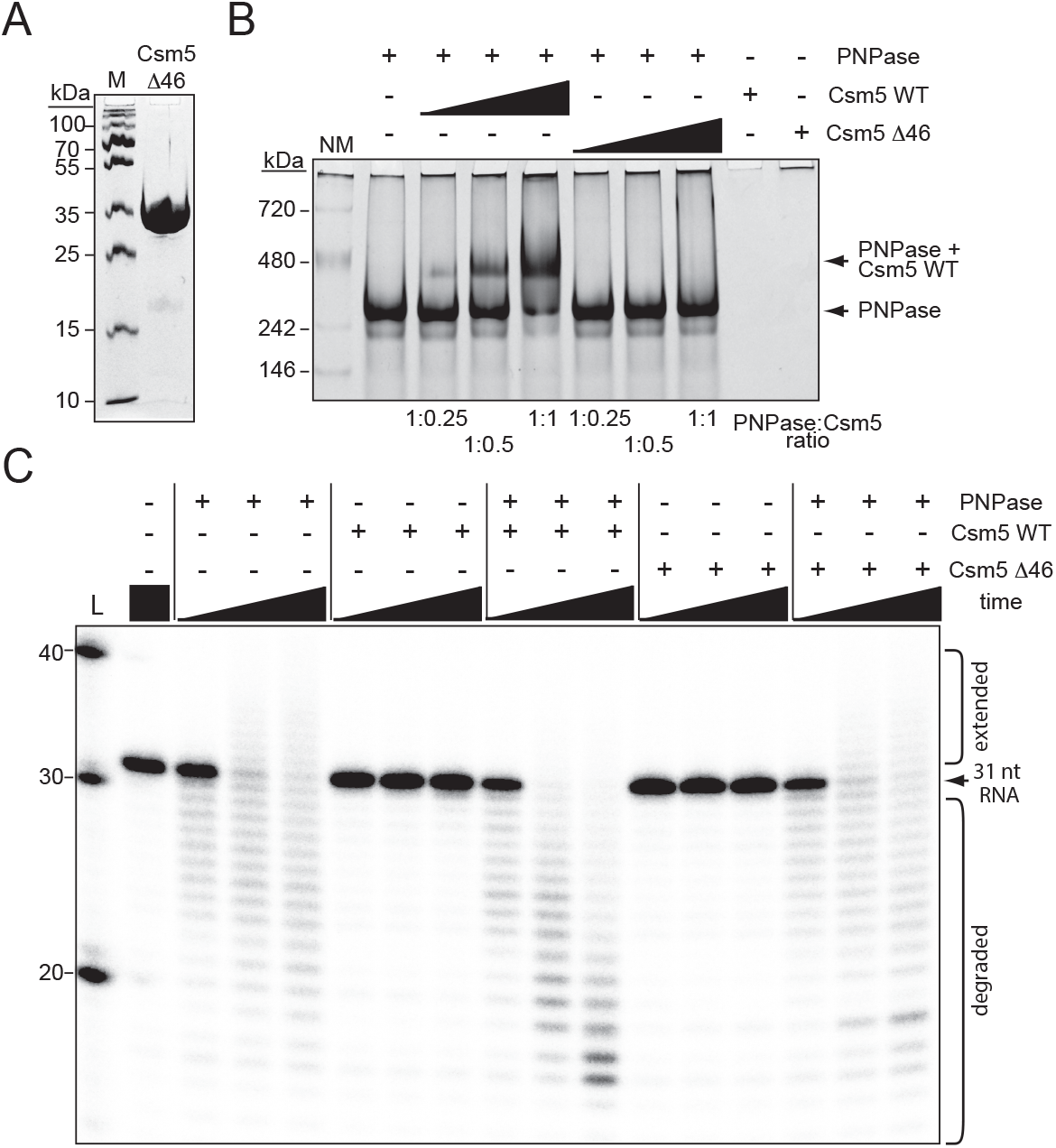
Csm5 interacts with and stimulates PNPase via a predicted disordered region. (A) Purified recombinant Csm5Δ46 is shown, in which IDR2 has been deleted. The protein was resolved on an SDS-PAGE gel and visualized using Coomassie G-250 staining. M, denaturing protein marker; kDa, kilodalton. See also Figure 5-source data 1. (B) PNPase was resolved on a native Cgel with increasing amounts of Csm5 (WT and Δ46). Shown is a representative of three independent trials. NM, native protein marker. See also Figure 5-source data 2. (C) Nuclease assays conducted with PNPase and/or Csm5 (WT and Δ46) are shown. In these assays, a 5’-end labelled 31-nucleotide RNA substrate was combined with indicated proteins, incubated at 37°C for increasing amounts of time (0.5, 5, and 15 minutes), and resolved on a denaturing gel. Shown is a representative of two independent trials. L, RNA Ladder. See also Figure 5-source data 3.

### RNase R and PNPase work synergistically to maintain robust anti-plasmid immunity

We next wondered about the extent to which RNase R and/or PNPase impact Type III CRISPR-Cas function. We began by testing immunity against diverse staphylococcal phages using various overexpression systems (Figure 6-figure supplement 1). First, immunity against siphovirus CNPx was tested in *S. epidermidis* LM1680 bearing p*crispr-cas/csm2*^*H6N*^ (Figure 6-figure supplement 1A). In this plasmid, the second spacer (*spc2*) targets the phage pre-neck appendage (*cn20*) gene (Daniel, Bonnen, and Fischetti 2007), which is likely to be expressed late in the phage infection cycle. Phage challenge assays were performed by spotting ten-fold dilutions of CNPx atop lawns of LM1680 cells bearing variants of the plasmid, incubating plates overnight, and enumerating phage plaques (i.e. clear zones of bacterial death) the next day. As expected, lawns of WT cells with the WT plasmid showed zero plaques, while lawns of WT cells bearing the empty vector allowed for tens of millions of plaques to form (measured as plaque-forming units per milliliter, pfu/ml) (Figure 6-figure supplement 1B). Interestingly, deletion of *rnr* alone or in combination with *pnp* caused no detectible defect in immunity. Surprisingly, even deletion of *csm5* from the plasmid had no impact on CRISPR function. In light of these observations, we wondered whether overexpressing the CRISPR-Cas system might compensate for mild defects. Thus, we tested another system that relies upon *S. epidermidis* RP62a (with an intact *crispr-cas* locus) bearing the multicopy plasmid p*crispr*, which contains a single repeat and spacer targeting phage(s) of interest (Bari et al. 2017). Since CNPx cannot form plaques on RP62a, phage challenge assays with podophage Andhra and myophage ISP were performed (Figure 6-figure supplement 1 C-F). Consistent with previous observations, the WT strain with the empty vector allows for formation of millions-billions of plaques, while RP62a strains with p*crispr* and appropriate phage-targeting spacers allow for zero plaques to form. Interestingly, RP62a strains devoid of *rnr* and/or *pnp* maintained robust immunity against both phages. Since previous work has shown that Type III-A immunity relies more heavily on the accessory nuclease Csm6 when phage late gene(s) are targeted owing to the lag in transcript/protospacer expression (Jiang, Samai, and Marraffini 2016), we explored the impact of targeting genes that are predicted to be expressed early vs. late in the infection cycle. Specifically, spacers targeting genes that encode Andhra’s DNA polymerase (early, *spcA1*), major tail protein (late, *spcA2*), and lysin-like peptidase (late, *spcA3*), as well as ISP’s lysin (late, *spcI*) were tested. Contrary to our expectations, robust anti-phage immunity was maintained in all strains. These results indicate that RNase R and PNPase may have little/no impact on immunity against diverse phages in these overexpression systems. We next tested anti-plasmid immunity using a conjugation assay that relies entirely upon chromosomally-encoded components (Fig. 6A and B). In this assay, *S. epidermidis* RP62a recipients are mated with *S. aureus* RN4220 cells harboring the conjugative plasmid pG0400. The first spacer in RP62a’s *crispr-cas* locus (*spc1)* bears complementarity to the nickase (*nes*) gene in pG0400 and therefore mitigates the conjugative transfer of the plasmid (Marraffini and Sontheimer 2008). Consistent with previous observations, mating assays performed with *S. aureus* RN4220/pG0400-WT donor cells and *S. epidermidis* RP62a WT recipients produced hundreds of transconjugants (i.e. *S. epidermidis* recipients that have acquired pG0400-WT); however, when *S. epidermidis* RP62a/*Δspc1-3* cells were used as recipients, >10,000 transconjugants were recovered (Fig. 6C). Interestingly, while RP62a/*Δpnp* performed similarly to the WT strain, RP62a/*Δrnr* exhibited a moderate attenuation in immunity, as evidenced by a significantly higher conjugation efficiency when compared to that of WT (Figure 6-source data 1). This defect was absent in the complemented strain (RP62a/*Δrnr::rnr\**), confirming that deletion of *rnr* is indeed responsible for the phenotype. Strikingly, the double-mutant (RP62a/*ΔrnrΔpnp*) showed a near complete loss of immunity, while the complemented strain RP62a/*ΔrnrΔpnp::rnr\** performed similarly to WT and RP62a/*Δpnp*. These data demonstrate that RNase R and PNPase work synergistically to promote anti-plasmid immunity.

**Figure 6.**
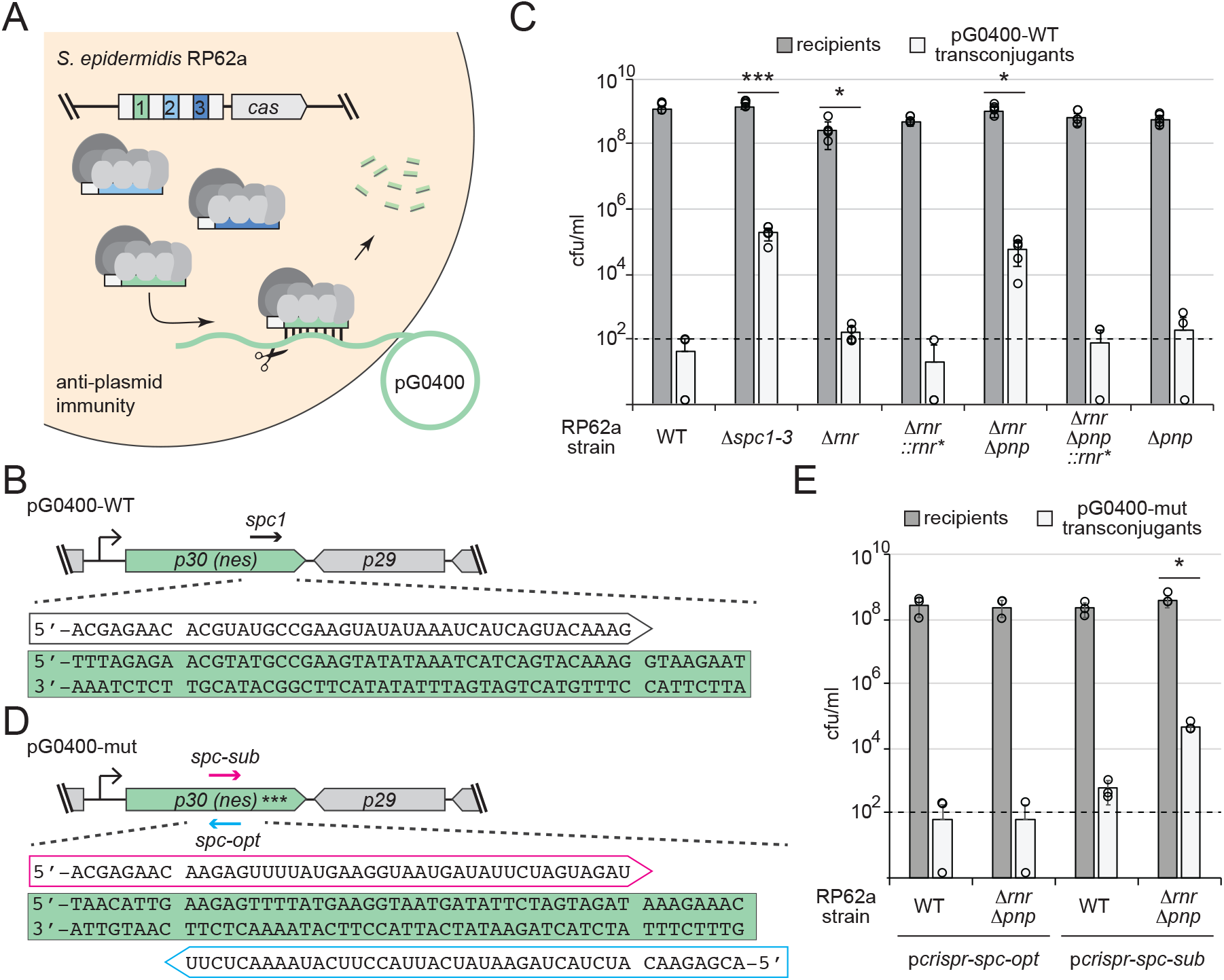
RNase R and PNPase work synergistically to promote robust anti-plasmid immunity. (A) Illustration of the anti-plasmid assay is shown in which the conjugative plasmid pG0400 is transferred from a *S. aureus* RN4220 donor (not shown) into various *S. epidermidis* RP62a recipient strains. The first spacer in the CRISPR locus (green square) bears complementarity to the nickase (*nes*) gene in pG0400. (B and D) Sequences of protospacers and corresponding crRNAs targeting pG0400-WT (B) and pG0400-mut (D). Protospacer sequences are highlighted in green, and targeting crRNA sequences are shown in unfilled arrows. In pG0400-mut, asterisks represent nine silent mutations in the *spc1* protospacer region. (C) Results from conjugation assays in which indicated *S. epidermidis* RP62a recipient strains were mated with *S. aureus* RN4220/pG0400-WT donor cells. See Figure 6-source data 1. (E) Results from conjugation assays in which various *S. epidermidis* RP62a recipient strains harboring indicated plasmids were mated with *S. aureus* RN4220/pG0400-mut donor cells. See Figure 6-source data 2. In panels C and E, numbers of recipients and transconjugants following mating are shown in cfu/ml (colony-forming units per mililiter). Graphs show an average of five (panel C) or three (panel E) independent trials (±S.D). Individual data points are shown with open circles, and data points on the *x*-axis represent at least one replicate where a value of 0 was obtained. The dotted line indicates the limit of detection for this assay. Two-tailed t-tests were performed on conjugation efficiencies to determine significance, and *p*-values of < 0.05 (*) and < 0.0005 (***) were obtained.

Although *spc1* was naturally-acquired in *S. epidermidis* RP62a and promotes anti-plasmid immunity in the WT background, it can be considered suboptimal because the crRNA that it encodes has the same (not complementary) sequence as the *nes* transcript (Fig. 6B). Accordingly, *spc1*-mediated anti-plasmid immunity was found to rely upon recognition of sparse antisense transcripts presumably originating from a weak promoter downstream of *nes* (Rostøl and Marraffini 2019). In light of these observations, we wondered whether the defect in anti-plasmid immunity in RP62a/*ΔrnrΔpnp* could be alleviated by targeting the more abundant *nes* transcript. To test this, we designed two spacers against the *nes* open reading frame, *spc-opt* and *spc-sub*, which encode crRNAs that are complementary to (optimal) and identical to (suboptimal) the *nes* transcript, respectively (Fig. 6D). Importantly, these spacers completely overlap and therefore share the same GC content. Further, the corresponding protospacers are devoid of complementarity between the 5’-tag on the crRNA and corresponding anti-tag region on the targeted transcript, a necessary condition that signals ‘non-self’ and licenses immunity (Marraffini and Sontheimer 2010). These spacers were inserted into p*crispr* to create p*crispr-spc-opt* and p*crispr-spc-sub*, and the plasmids were introduced into RP62a WT and RP62a/*ΔrnrΔpnp*. In order to eliminate the effects of the chromosomally-encoded *spc1* in these strains, conjugation assays were performed with *S. aureus* RN4220 cells bearing pG0400-mut, a variant of pG0400-WT that is not recognized by the *spc1* crRNA owing to the presence of nine silent mutations across the protospacer (Marraffini and Sontheimer 2008). We found that similarly to *spc1, spc-sub* mediates anti-plasmid immunity in RP62a WT, but fails to do so in RP62a/*ΔrnrΔpnp* (Fig. 6E). In contrast, *spc-opt* facilitates robust immunity in both WT and the double-mutant. Altogether, these data support the notion that the nuclease activities of RNase R and PNPase are essential to maintain robust immunity when targeted transcripts are in low abundance.

## Discussion

Here, we elucidate the complete pathway for crRNA maturation in a model Type III-A CRISPR-Cas system and expand the repertoire of known accessory nucleases required for immunity (Fig. 7). Most functionally-characterized Type III systems generate mature crRNAs that vary in length on their 3’-ends by six nucleotide increments (Hale et al. 2009; Hatoum-Aslan, Maniv, and Marraffini 2011; Staals et al. 2013; Tamulaitis et al. 2014), and while this periodic cleavage pattern is known to derive from the protection offered by variable copies of Csm3/Cmr4 subunits within effector complexes (Hatoum-Aslan et al. 2013; You et al. 2019; Osawa et al. 2015; Dorsey, Huang, and Mondragon 2019), the identity of the nuclease(s) responsible for crRNA 3’-end maturation and the functional significance of this additional processing step have long remained a mystery. Here, we demonstrate that two housekeeping nucleases, RNase R and PNPase, work in concert to trim the 3’-ends of intermediate crRNAs (Fig. 1 and 2) and promote robust anti-plasmid immunity in *S. epidermidis* (Fig. 6). Since intermediate crRNAs can mediate a successful immune response under certain conditions, such as when the CRISPR-Cas system is overexpressed (Figure 6-figure supplement 1) or when a highly-expressed transcript is targeted (Fig. 6 D and E), the functional value of crRNA maturation is likely nominal and may occur simply as a consequence of the pre-emptive recruitment of RNase R and PNPase to the effector complex to assist during interference.

**Figure 7.**
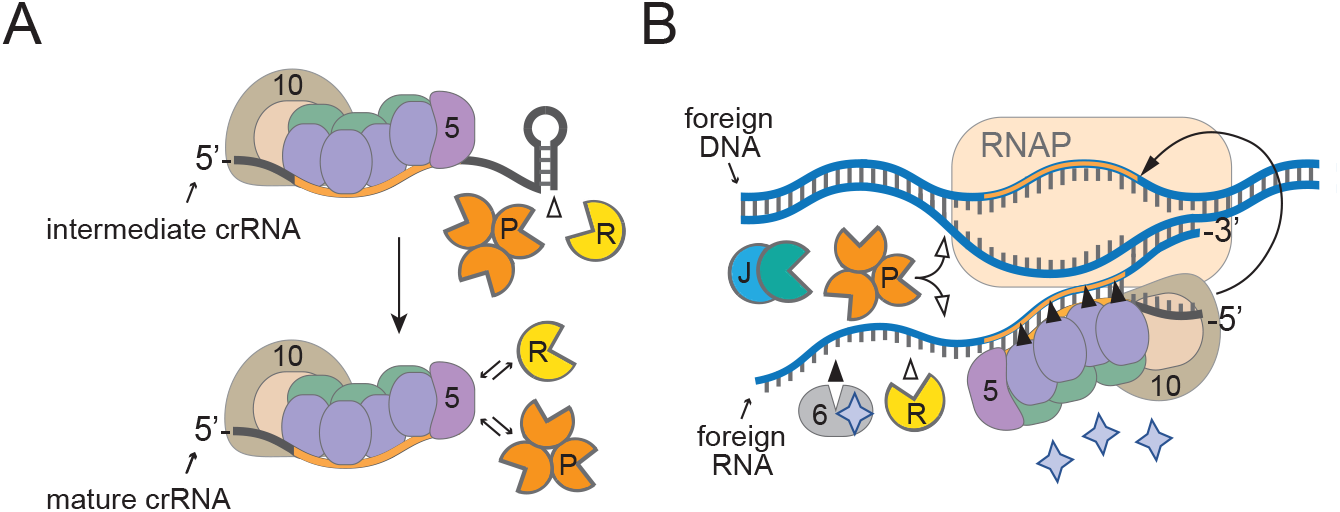
A model for how diverse housekeeping nucleases are enlisted to ensure successful defense. (A) During Cas10-Csm complex assembly, Csm5 recruits and/or stimulates RNase R and PNPase through direct interactions. The unprotected 3’-ends of intermediate crRNAs are trimmed as a consequence of nuclease recruitment, resulting in the generation of the shorter mature species. (B) During interference, RNase R and PNPase work synergistically to help degrade invading nucleic acids alongside other Cas and non-Cas nucleases. Filled triangles illustrate events catalyzed by Cas proteins, and open triangles illustrate events catalyzed by non-Cas nucleases. Purple stars represent cyclic oligoadenylate molecules produced by Cas10. 5, Csm5; 6, Csm6; 10, Cas10; R, RNase R; P, PNPase; J, RNase J1/J2; RNAP, RNA Polymerase.

It is now well understood that most, if not all, Type III CRISPR-Cas systems rely upon RNA recognition to eliminate invading DNA (Samai et al. 2015; Kazlauskiene et al. 2016; Estrella, Kuo, and Bailey 2016; Elmore et al. 2016; Liu, Iavarone, and Doudna 2017), and this feature presents unique challenges owing to the fact that targeted transcripts can have variable levels of expression. Previous studies have shown that while highly-expressed target RNAs elicit a robust and sustained immune response that results in the swift elimination of nucleic acid invaders, low-abundance targets evoke a weak immune response, and corresponding invaders are more difficult to clear (Goldberg et al. 2014; Jiang, Samai, and Marraffini 2016; Rostøl and Marraffini 2019). In the latter scenario, the Cas10-Csm complex requires the help of Csm6, an accessory nuclease encoded in the CRISPR-Cas locus that is not part of the complex. Csm6 has nonspecific endoribonuclease activity which is stimulated when bound to cOAs produced by Cas10 (Kazlauskiene et al. 2017; Niewoehner et al. 2017; Nasef et al. 2019), and Csm6-mediated degradation of transcripts derived from both the invader and host causes growth arrest until the foreign nucleic acids are cleared (Rostøl and Marraffini 2019). Interestingly, Csm6 is dispensable for immunity when targeted transcripts are highly-expressed (Jiang, Samai, and Marraffini 2016; Rostøl and Marraffini 2019), similar to RNase R and PNPase (Fig. 6). Altogether, our observations support a model in which RNase R and PNPase are recruited as accessory nucleases to ensure a successful defense against nucleic acid invaders, particularly when targeted transcripts have low abundance (Fig. 7).

In spite of these similarities with Csm6, RNase R and PNPase have distinct functional roles in the cell and mechanisms by which they are enlisted for defense. Unlike Csm6, PNPase and RNase R are 3’-5’ exonucleases primarily involved in housekeeping functions--PNPase is a member of the RNA degradosome, a multi-enzyme complex that catalyzes RNA processing and degradation, and RNase R performs similar/overlapping functions, but works independently of the degradosome in most organisms (Bechhofer and Deutscher 2019; Tejada-Arranz, de Crecy-Lagard, and de Reuse 2020). In addition to its RNase activity, PNPase has the capacity to degrade single-stranded DNA (Walker et al. 2017), presumably to facilitate DNA repair (Cardenas et al. 2009). These activities are harnessed by the Cas10-Csm complex through direct interactions—Csm5 essentially borrows both nucleases through weak/transient interactions, and PNPase’s nuclease activity is further stimulated when bound to Csm5 (Figs. 3 and 5). While the specific binding site for RNase R remains unknown, a predicted IDR on the C-terminus of Csm5 is responsible for recruitment and stimulation of PNPase (Figs. 4, 5, Figure 4-figure supplement 1, and Figure 5-figure supplement 1). Since RNase R and PNPase are each about twice the size of Csm5, their association with the Cas10-Csm complex is likely mutually-exclusive, and their docking site(s) may also overlap with other subunits within the complex. Regardless of the precise molecular requirements for recruitment, Cas10-Csm’s ability to interface with diverse cellular nucleases is remarkable and bears striking parallels to the assembly of prokaryotic degradosomes and eukaryotic granules, which rely upon “hub” proteins to recruit multiple members of enzyme complexes through transient interactions with one or more IDRs (Tejada-Arranz, de Crecy-Lagard, and de Reuse 2020). Given that most Type III CRISPR-Cas systems possess Csm5 homologs (Makarova, Wolf, et al. 2020), and RNase R and PNPase are evolutionarily conserved in prokaryotes and eukaryotes (Zuo and Deutscher 2001), their enlistment in Type III CRISPR-Cas defense may be a common feature in diverse organisms.

### Ideas and Speculation

Harnessing the activities of housekeeping nucleases and channeling their diverse activities towards defense may have evolved as a strategy to minimize the genetic footprint of complex immune systems while cutting the energetic costs associated with manufacturing enzymes with redundant functions. Supporting this notion, diverse CRISPR-Cas systems have been shown to tap into the pool of cellular housekeeping nucleases to perform different steps in their immunity pathways. The earliest example of this phenomenon was discovered over a decade ago in the Type II CRISPR-Cas system of *Streptococcus pyogenes*, in which processing of both crRNAs and the trans-encoded small RNA (tracrRNA) was shown to rely upon RNase III (Deltcheva et al. 2011). RNase III-mediated crRNA/tracrRNA processing is now considered a universal feature of Type II systems (Makarova, Wolf, et al. 2020). In addition, new spacer acquisition (i.e. adaptation) in Type I and II systems has been shown to rely upon the DNA repair machinery RecBCD and AddAB in gram -negative and -positive organisms, respectively (Levy et al. 2015; Modell, Jiang, and Marraffini 2017). Also, we showed that RNase J2 plays a critical role in interference in the Type III-A system of *S. epidermidis* (Chou-Zheng and Hatoum-Aslan 2019). Beyond these more common systems, CRISPR-Cas variants in which one or more *cas* nucleases are missing were shown to rely upon degradosome nucleases to perform essential functionalities. In one such example, a Type III-B variant in *Synechocystis* 6803 which lacks a Cas6 homolog relies upon RNase E to catalyze processing of pre-crRNAs (Behler et al. 2018). In another more extreme example, a unique CRISPR element in *Listeria monocytogenes* which is completely devoid of *cas* genes was shown to utilize PNPase for crRNA processing and interference (Sesto et al. 2014). Since housekeeping nucleases are needed on a daily basis and therefore less likely to be lost via natural selection, it is plausible that their enlistment in nucleic acid defense may be more widespread than currently appreciated.

## Materials and Methods

### Bacterial strains, phages, and growth conditions

*S. aureu*s RN4220 was propagated in Tryptic Soy Broth (TSB) medium (BD Diagnostics, NJ, USA). *S. epidermidis* LM1680 and RP62a were propagated in Brain Heart Infusion (BHI) medium (BD Diagnostics, NJ, USA). *E. coli* DH5α was propagated in Luria Bertani (LB) broth (VWR, PA, USA), and *E. coli* BL21 (DE3) was propagated in Terrific broth (TB) medium (VWR, PA, USA) for protein purification.

Corresponding media was supplemented with the following: 10 µg/ml chloramphenicol (to select for p*crispr-spc*-, p*crispr-cas*-, and pKOR1-based plasmids), 15 µg/ml neomycin (to select for *S. epidermidis* cells), 5 µg/ml mupirocin (to select for pG0400-based plasmids), 50 µg/ml kanamycin (to select for pET28b-His10Smt3-based plasmids), and 30 µg/ml chloramphenicol (to select for *E. coli* BL21 (DE3)). Phages CNPx and ISP were propagated using *S. epidermidis* LM1680 as host, and phage Andhra was propagated using *S. epidermidis* RP62a as host. For phage propagation, overnight cultures of the corresponding hosts were diluted at 1:100 in BHI supplemented with 5 mM CaCl_2_ and phage (10^6^-10^8^ pfu/ml). Cultures were incubated at 37°C with agitation for 5 hr. One-fifth the volume of host cells (grown to mid-log) was added into the bacteria-phage mixture and incubated for an additional 2 hr at 37°C with agitation. Cells were pelleted at 5000 x *g* for 5 min at 4°C, and the supernatant containing phage was filtered using a 0.45 µm syringe filter. Phage titers were determined using the double-agar overlay method as described in (Cater et al. 2017). Briefly, a semisolid layer of 0.5 X heart infusion agar (HIA) medium (Hardy Diagnostics, CA, USA) containing 5 mM CaCl_2_ and a 1:100 dilution of overnight host culture was overlaid atop a solid layer of Tryptic Soy Agar (TSA) (BD Diagnostics, NJ, USA) plates supplemented with 5 mM CaCl_2_. Filtered phages were diluted in ten-fold dilutions and spotted atop the semisolid layer, spots were air-dried, and plates were incubated overnight at 37°C. Plaques were then enumerated, and phage titers in plaque-forming units/ml (pfu/ml) were determined.

### Construction of pKOR1-based plasmids and transformation into S. epidermidis LM1680

The pKOR1 system (Bae and Schneewind 2006) was used to create in-frame deletions of *rnr* (encodes RNase R) and to re-insert an *rnr* variant (*rnr*\*, which has two silent mutations, see Figure 1-figure supplement 1) into *S. epidermidis* LM1680 WT and Δ*pnp* strains, and RP62a WT and Δ*pnp* strains. The pKOR1 vector was used as a template to amplify the backbone for all pKOR1-based constructs with primers A481/L138 via PCR amplification. The plasmid, pKOR1-Δ*rnr*, was created using a 3-piece Gibson assembly (Gibson et al. 2009) and used to delete *rnr*. Briefly, two DNA fragments flanking upstream and downstream of *rnr* were obtained via PCR amplification using primers L139/L140 and L141/L142 (Table S2), respectively, and *S. epidermidis* RP62a WT as template. The PCR products of these flanks and pKOR1 backbone were purified using the EZNA Cycle Pure Kit (Omega Bio-tek, CA, USA) and Gibson assembled. The plasmid pKOR1-*rnr*\* was created via a 3-piece Gibson assembly and used to re-introduce *rnr*\* back to Δ*rnr* strains as follows. Briefly, primers L154/L155 (which bind to *rnr*) were designed to introduce two silent mutations which remove a native EcoR1 restriction site (Figure 1-figure supplement 1 C and D and Table S2). Then, upstream and downstream flanking regions of *rnr* were amplified with PCR using primers L139/L155 and L154/L142, respectively, with *S. epidermidis* RP62a WT as template. PCR products of these flanks and pKOR1 backbone were purified using the EZNA Cycle Pure Kit and Gibson assembled. All assembled constructs were transformed via electroporation into *S. aureus* RN4220. Four transformants were selected for each construct and the presence of the plasmid was confirmed using PCR amplification and DNA sequencing with primers L145/L146 (Table S2). Confirmed plasmids were extracted using the EZNA Plasmid DNA Mini Kit (Omega Bio-tek, CA, USA) and introduced into *S. epidermidis* LM1680 WT and Δ*pnp* via electroporation. Plates were incubated at 30°C for 48 hr. Four transformants were selected and analyzed using PCR amplification and DNA sequencing with primers L145/L146 to confirm presence of plasmid. Confirmed *S. epidermidis* LM1680 WT and Δ*pnp* transformants were used to proceed with mutagenesis and *S. epidermidis* LM1680 WT harboring appropriate plasmid was used to transfer the pKOR1-based plasmids into *S. epidermidis* RP62a WT and Δ*pnp* using phage-mediated transduction.

### Transduction of pKOR1-based plasmids into S. epidermidis RP62a

The temperate phage CNPx was used to transduce pKOR1-based plasmids from *S. epidermidis* LM1680 WT into *S. epidermidis* RP62a WT and Δ*pnp* as described previously in (Chou-Zheng and Hatoum-Aslan 2019) with slight modifications. Briefly, overnight cultures of *S. epidermidis* LM1680 WT and Δ*pnp* strains harboring pKOR1-based plasmids were used to propagate phage CNPx as described above. Bacteria-phage cultures were incubated at 37°C with agitation for 5 hr, or until cell lysis. Cells were pelleted at 5000 x *g* for 5 min at 4°C, and the phage lysates were passed through a 0.45 µm syringe filter. Filtered phage lysates were then mixed with mid-log *S. epidermidis* RP62a cells in a 1:10 dilution and incubated at 37°C for 20 min. Bacteria-phage cultures were pelleted at 5000 x *g* for 1 min. Cell pellets were washed twice with 1 mL of fresh BHI, and the final pellets were resuspended in 200 µl of fresh BHI and plated entirety onto BHI agar containing appropriate antibiotics. Plates were then incubated at 30°C for 48 hr. Four transductants were selected for each construct and the plasmid’s presence was confirmed using PCR amplification and DNA sequencing with primers L145/L146 (Table S2).

### Generation of S. epidermidis Δrnr and ΔrnrΔpnp

*S. epidermidis* strains bearing pKOR1-*Δrnr* and pKOR1-*rnr*\* were used to generate all corresponding mutants using allelic replacement (Bae and Schneewind 2006) as described previously in (Chou-Zheng and Hatoum-Aslan 2019). Four independent *Δrnr* and *ΔrnrΔpnp* deletion strains (i.e. biological replicates) were created and confirmed using PCR amplification and DNA sequencing with primers L143/L157. Four independent *Δrnr::rnr\** and *ΔrnrΔpnp::rnr\** complemented strains (i.e. biological replicates) were created and confirmed using three methods: PCR amplification, DNA sequencing with primers L143/L153, and EcoRI (New England Biolabs, MA, USA) digestion of PCR products (Table S2).

### Construction of pcrispr-cas/csm5^H6N^ Δ18, Δ31, and Δ46

All p*crispr-cas*-based plasmids were constructed using a 3-piece Gibson assembly. The p*crispr-cas* plasmid (Hatoum-Aslan et al. 2013) was used as a template to amplify the backbone for these constructs. The three PCR products for p*crispr-cas/csm5*^*H6N*^Δ18 were obtained using primer sets F063/F066, F067/L247, and F062/L246 (Table S2). The three PCR products for p*crispr-cas/csm5*^*H6N*^Δ31 were obtained using primer sets F061/F066, F067/L265, and F046/L264. The three PCR products for p*crispr-cas/csm5*^H6N^Δ46 were obtained using primer sets F061/F066, F067/L275, and F046/L274. All PCR products were purified using the EZNA Cycle Pure Kit and Gibson assembled. All assembled constructs were introduced into *S. aureus* RN4220 via electroporation. Four transformants were selected for each construct and confirmed to harbor the plasmid using PCR amplification and DNA sequencing with primers A416/F113. Confirmed constructs were extracted using the EZNA Plasmid DNA Mini Kit and transferred via electroporation into *S. epidermidis* LM1680 WT. Four transformants were selected and analyzed with PCR amplification and DNA sequencing using primers A416/F113 to confirm the presence of desired plasmids.

### Construction of pcrispr-spc-based plasmids

Spacers were designed using the protospacer selector tool (https://github.com/ahatoum/CRISPRCas10-Protospacer-Selector) described in (Bari et al. 2017). Briefly, spacers were designed to target specific gene region of the corresponding phage, or the *nes* gene of conjugative plasmid pG0400, that bear no complementarity between the eight-nucleotide tag on the 5′-end of the crRNA (5′-ACGAGAAC), and the “anti-tag” region adjacent to the protospacer. Selected spacers were introduced into the template p*crispr-spcA1* (referred to as p*crispr-spcA2* in (Bari et al. 2017)) via inverse PCR using the primers listed in Table S2. All PCR products were purified using the EZNA Cycle Pure Kit. Linear products were phosphorylated with T4 Polynucleotide Kinase (New England Biolabs, MA, USA) for 1 hr at 37°C and circularized with T4 DNA Ligase (New England Biolabs, MA, USA) overnight at room temperature using buffers and instructions provided by the manufacturer. All assembled constructs were introduced into *S. aureus* RN4220 via electroporation. Four transformants were selected for each construct and confirmed via PCR amplification and DNA sequencing with primers A200/F052. Confirmed plasmids were extracted using the EZNA Plasmid DNA Mini Kit and introduced into *S. epidermidis* RP62a via electroporation. Four transformants were selected and analyzed with PCR amplification and DNA sequencing with primers A200/F052 to confirm presence of desired plasmids.

### *CRISPR-Cas10 functional* assays

Conjugation assays using pG0400-WT and pG0400-mut were carried out by filter mating *S. aureus* donor strains harboring these plasmids with various *S. epidermidis* recipient strains as described previously in (Walker and Hatoum-Aslan 2017). The conjugation data reported represents mean values (±S.D.) of three to five independent trials (see appropriate figure legends for details). Phage challenge assays were carried out by spotting ten-fold dilutions of phages atop lawns of cells as previously described in (Chou-Zheng and Hatoum-Aslan 2019). Phage CNPx was used to infect *S. epidermidis* LM1680 WT and mutant strains carrying p*crispr-cas*-based plasmids. Phages Andhra and ISP were used to infect *S. epidermidis* RP62a WT and mutant strains carrying p*crispr-spc*-based plasmids. The phage challenge data reported represents mean values (±S.D.) of three independent trials.

### Purification of Cas10-Csm complexes from S. epidermidis LM1680

Cas10-Csm complexes containing a 6-His tag on the N-terminus of Csm2 or Csm5 in pcrispr-cas-based plasmids were overexpressed in *S. epidermidis* LM1680 cells, harvested and stored exactly as described in (Chou-Zheng and Hatoum-Aslan 2017). Final pellets were purified following the first affinity chromatography protocol (Ni^2+^-affinity chromatography) with slight modifications. Briefly, cell pellets were resuspended in 10 ml of lysis buffer A (22 mM MgCl_2_, 44 μg/ml lysostaphin) and incubated in a water bath at 37°C for 1 hr. Lysed cells were then resuspended with 10 ml of lysis buffer B (50 mM NaH_2_PO_4_, 300 mM NaCl, pH 8.0) supplemented with 20 mM imidazole, 0.1% Triton X-100, and one complete EDTA-free protease inhibitor tablet (Roche, Basel, Switzerland). Cells were homogenized by inverting the tube several times until the mixture becomes very viscous (a 5 min incubation period at room temperature might be necessary to achieve viscosity). Cells were then sonicated, and insoluble material was removed via centrifugation and filtration. Cleared lysates were passed through a 5 ml gravity column (G-Biosciences, MO, USA) containing 1.5 ml of Ni^2+^-NTA agarose resin (ThermoFisher Scientific, MA, USA) pre-equilibrated with lysis buffer B. Nickel-bound complexes were then washed with 15 ml of lysis buffer B supplemented with 20 mM imidazole and 5% glycerol, followed by another 15 ml wash of lysis buffer B supplemented with 20 mM imidazole and 10% glycerol. Complexes were then eluted with five 600 µL aliquots each of lysis buffer B supplemented with 250 mM imidazole and 10% glycerol. Complexes were resolved on a 15% SDS PAGE and visualized with Coomassie G-250. A pre-stained protein standard (New England Biolabs, MA, USA) was used to estimate molecular weight. Protein concentrations were determined using absorbance measurements at 280 nm (A280) with a NanoDrop2000 spectrophotometer (ThermoFisher Scientific, MA, USA).

### Reconstitution of crRNA maturation

300 pmol of Cas10-Csm complexes purified from LM1680/*ΔrnrΔpnp* were combined with 100 pmol of purified Cbf1, PNPase, and/or RNase R in Nuclease Buffer A (25 mM Tris-HCl pH 7.5, 2 mM DTT) supplemented with 10 mM MgCl_2_ (PNPase and RNase R), or 10 mM MnCl_2_ (Cbf1). Nuclease reactions were carried out at 37°C for 30 min, or for a time course of 15, 30, and 60 min. Reactions were halted on ice for 10 min, and then crRNAs were extracted and visualized.

### Extraction and visualization of crRNAs

Total crRNAs were extracted from purified Cas10-Csm complexes as described previously in (Chou-Zheng and Hatoum-Aslan 2019) with slight modifications. Briefly, 300 to 600 pmols of purified complexes were resuspended in 750 µL TRIzol Reagent (Invitrogen, NY, USA) and subsequent RNA extraction steps were completed as recommended by the manufacturer. Extracted crRNAs were end-labeled with T4 Polynucleotide Kinase in a reaction containing γ-[^32^P]-ATP (PerkinElmer, MA, USA), and resolved on an 8% Urea PAGE. The gel was exposed to a storage phosphor screen and visualized using an Amersham Typhoon biomolecular imager (Cytiva, MA, USA). For densitometric analysis, the ImageQuant software was used. Percent of intermediate crRNAs was obtained using the following equation: [intensity of intermediate crRNA signal (71 nt) ÷ sum of signal intensities for the dominant crRNA species (71 nt + 43 nt + 37 nt + 31 nt)] × 100%. The data reported represents mean values (±S.D.) of two to four independent trials (see appropriate figure legends for details).

### Construction of pET28b-His_10_Smt3-based plasmids

pET28b-His_10_Smt3-*csm5Δ46* was constructed via inverse PCR using primers L274/L275 (Table S2) and template pET28b-His_10_Smt3-*csm5* (Walker et al. 2017). PCR products were digested with DpnI (New England Biolabs, MA, USA) as indicated by the manufacturer and purified using the EZNA Cycle Pure Kit. Purified PCR products were then 5’-phosphorylated with T4 Polynucleotide Kinase for 1 hr at 37°C and circularized with T4 DNA Ligase overnight at room temperature using buffers and instructions provided by the manufacturer. Ligated pET28b-His_10_Smt3-*csm5Δ46* constructs were introduced into *E. coli* DH5α via chemical transformation. Three transformants were selected, screened, and confirmed using PCR amplification and DNA sequencing with primers T7P/T7T. Confirmed plasmids were purified using the EZNA Plasmid DNA Mini Kit and introduced into *E. coli* BL21 (DE3) via chemical transformation for protein purification. Three transformants were selected and re-confirmed using PCR amplification and DNA sequencing with primers T7P/T7T (Table S2).

### Overexpression and purification of recombinant Csm5, Csm5Δ46, PNPase, RNase R, and Cbf1 from E. Coli

*E. coli* BL21 (DE3) cells bearig pET28b-His_10_Smt3-based plasmids were grown, induced, and recombinant proteins purified as previously described (Walker et al. 2017) with slight modifications. Following cell harvesting, pellets were placed on ice and resuspended in 30 ml of Buffer A (50 mM Tris-HCl pH 6.8, 1.25 M NaCl, 200 mM Li_2_SO4, 10% sucrose, 25 mM Imidazole) supplemented with one complete EDTA-free protease inhibitor tablet (Roche, Basel, Switzerland), 0.1 mg/ml lysozyme, and 0.1% Triton X-100. Cells were incubated for 1 hr at 4°C with constant rotation, then sonicated. Insoluble material was removed via centrifugation and filtration. Cleared lysates were mixed with 4 ml of Ni^2+^-NTA agarose resin pre-equilibrated with Buffer A, then mixed for 1 hr at 4°C with constant rotation. The resin was pelleted, washed with 40 ml of Buffer A, and pelleted again. The resin was then resuspended in 5 ml of Buffer A and transferred to a 5 ml gravity column. The resin was further washed with 20 ml of Buffer A. Proteins were eluted stepwise with three aliquots of 1 ml each of IMAC buffer (50 mM Tris-HCl pH 6.8, 250 mM NaCl, 10% glycerol) containing 50, 100, 200 and 500 mM imidazole. Eluted protein fractions were resolved on a 15% SDS PAGE, visualized with Coomassie G-250, and estimated molecular weight was determined with pre-stained protein standard. Fractions containing the desired protein were pooled and mixed with SUMO Protease (MCLAB, CA, USA) with the provided SUMO buffer (salt-free). The mixtures were dialyzed for 3 hr against IMAC buffer containing 25 mM imidazole. The dialysate was mixed with 2 ml of Ni^2+^-NTA agarose resin (pre-equilibrated with IMAC buffer containing 25 mM imidazole) and mixed for 1 hr with constant rotation at 4°C. The digested mixture was passed through a 5 ml gravity column, and the untagged protein was collected in the flow-through. Additional untagged protein was collected by flowing through the column two 1 mL aliquots each of IMAC buffer containing 50, 100 and 500 mM imidazole. Proteins were resolved, visualized, and estimated as described above. Protein concentrations less than 1 mg/mL were concentrated using a 10K MWCO centrifugal filter (Pall Corporation, NY, USA). Protein concentrations were determined using absorbance measurements at 280 nm (A280) with a NanoDrop2000 spectrophotometer.

### Nuclease assays

A 31-nt Single stranded RNA substrate (5’ ACGAGAACACGUAUGCCGAAGUAUAUAAAUC) was 5’ end-labeled with T4 Polynucleotide Kinase and γ-[32P]-ATP, and purified with a G25 column (IBI Scientific, IA, USA). Labeled substrate was incubated with 1 pmol of PNPase and 5 pmol of Csm5 WT or Csm5Δ46 in Nuclease Buffer B (25 mM Tris-HCl pH 7.5, 2 mM DTT, 10 mM MgCl2). Nuclease reactions were carried out at 37°C in a time course of 0.5, 5, and 15 min, and quenched by adding an equal volume of 95% formamide loading buffer. Reactions were resolved on a 15% UREA PAGE. Gel was exposed to a storage phosphor screen and visualized using an Amersham Typhoon biomolecular imager.

### Native gel electrophoresis

All native electrophoresis gels were run in a Tetra Vertical Electrophoresis Chamber (Mini-PROTEAN® Tetra Cell, Bio-Rad, CA, USA). Recombinant PNPase (100 pmols) alone or in combination with Csm5 WT, or Csm5Δ46, (25, 50, and 100 pmols) was resolved in a 6% native polyacrylamide gel (29:1 acrylamide/bisacrylamide) of 0.75 mm thick. Tris-glycine buffer (25 mM Tris, 250 mM glycine, pH 8.5) was used to prepare and run the gels. Recombinant RNase R (30 pmols) or Bovine Serum Albumin (BSA, Sigma-Aldrich, MO, USA) were alone or combined with Csm5 WT or Csm5Δ46 (90, 180, 225, and 270 pmols) were resolved in 6% native polyacrylamide gels (29:1 acrylamide/bisacrylamide) of 1.0 mm thick. Tris-CAPS buffer (60 mM Tris, 40 mM CAPS, pH 9.3-9.6) was used to prepare and run the gels. Native gel electrophoresis was conducted on an ice-water bath for 100-130 min at 90 V. Proteins were visualized with Coomassie G-250. NativeMark Protein Standard (Thermo Fisher Scientific) was used to estimate molecular weight.

### Statistical analyses and replicate definitions

All graphed data represents the mean (± S.D.) of *n* replicates, where the *n* value is indicated in figure legends and source data files. Average values were analyzed in pairwise comparisons using two-tailed t-tests, and *p* values below 0.05 were considered statistically significant. Sample sizes were empirically determined, and no outlyers were observed or omitted. The following terms are used to describe the types of repetitions where appropriate in figure legends and source data files: independent trials, independent transformants, and biological replicates. Independent trials refer to repetitions of the same experiment conducted at different times; independent transformants refer to one bacterial strain into which a construct was independently introduced; and biological replicates refer to different bacterial mutants that were independently created.

### Materials Availability

All bacterial strains and constructs generated in this study can be made available by the corresponding author (A. H-A.) upon written request.

## Code Availability

There was no new code generated in this work; however, a previously-generated code was re-used to identify permissive protospacers in the *nes* gene for Type III-A CRISPR-Cas immunity (in Fig. 6D and E). For this experiment, spacers were designed using the publicly available protospacer selector tool (https://github.com/ahatoum/CRISPRCas10-Protospacer-Selector) described in (Bari et al. 2017)

## Acknowledgements

We would like to acknowledge funding for this project from the National Science Foundation CAREER award (MCB/2054755). A.H-A. is also funded by an Investigators in the Pathogenesis of Infectious Disease award from the Burroughs Wellcome Fund.

## Supplementary Figures and Tables

**Figure 1-figure supplement 1.**
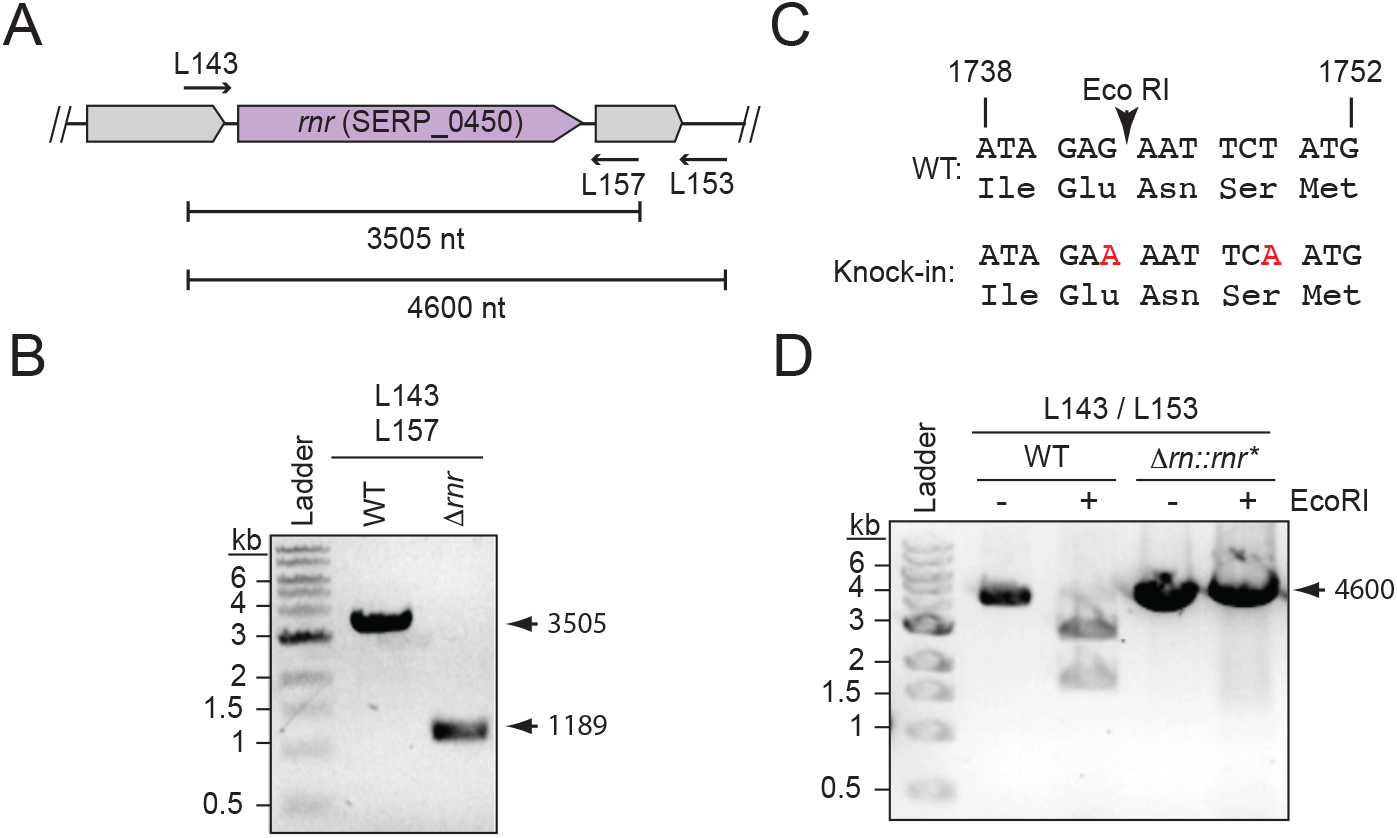
Confirmation of *rnr* knock-out and knock-in *S. epidermidis* strains. (A) Illustration of the *rnr* genomic locus and corresponding primers used to amplify the gene (see Table S2 for primer sequences). (B) PCR products of *rnr* region from representative *S. epidermidis* WT and knock-out strains are shown. The *rnr* gene was deleted using the pKOR1 allelic replacement system (Bae and Schneewind 2006). Following mutagenesis, the *rnr* deletion was confirmed by PCR amplification of the genomic locus with primers L143/L157 and sequencing the PCR product. See Figure 1-figure supplement 1-source data 1. (C) A segment of the coding region of *rnr* into which two silent mutations were introduced (red nucleotides) to distinguish the knock-in strain from wild-type. The two silent mutations in *rnr* gene (*rnr*\*) remove a native EcoRI restriction site naturally found within *rnr* gene. The *rnr*\* variant was replaced into its original genetic locus in knockout strains using the pKOR1 allelic replacement system (Bae and Schneewind 2006). (D) PCR products of *rnr* region from representative *S. epidermidis* wild-type and knock-in strains are shown. Following mutagenesis, *rnr*\* was confirmed by PCR amplifying genomic DNA with primers L143/L153, and digesting PCR products with EcoRI restriction enzyme, and sequencing the PCR product. kb, kilobase. See Figure 1-figure supplement 1-source data 2.

**Figure 2-figure supplement 1.**
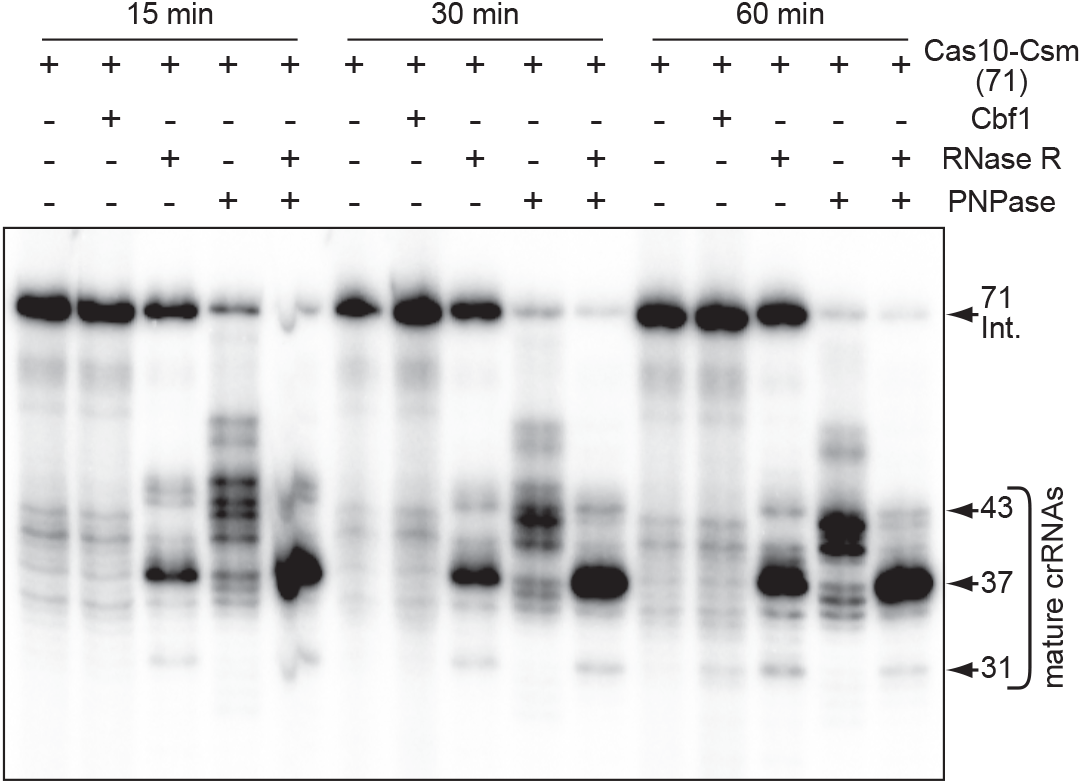
RNase R alone cannot complete crRNA maturation in a purified system. Total crRNAs associated with purified Cas10-Csm (71) complexes are shown after digestion with indicated nucleases for different timepoints (15, 30, and 60 minutes). CrRNAs were extracted using trizol reagent, radiolabeled at their 5’-ends, and resolved on a denaturing gel. The gel shown is a representative of two independent trials. See Figure 2-figure supplement 1-source data 1.

**Figure 3-figure supplement 1.**
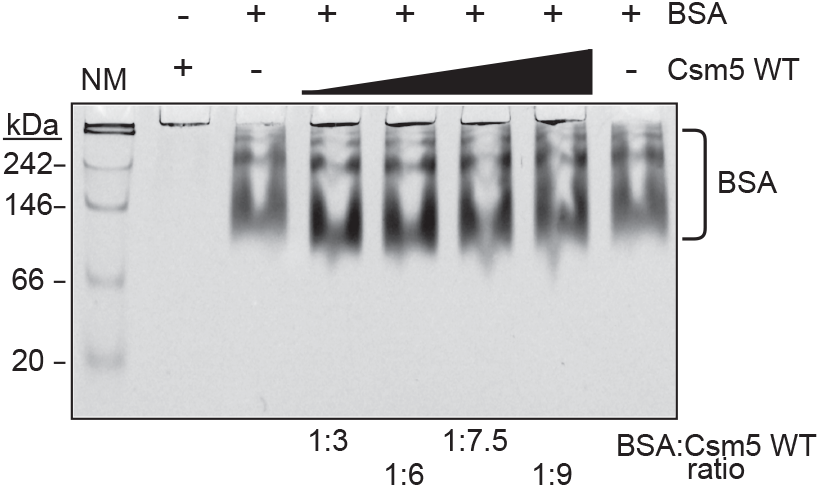
Csm5 does not interact with BSA (bovine serum albumin). Shown is a native gel in which BSA was resolved alone or with increasing concentrations of Csm5 WT. The gel shown is a representative of three independent trials. NM, native protein marker; kDa, kilodalton. See Figure 3-figure supplement 1-source data 1.

**Figure 4-figure supplement 1.**
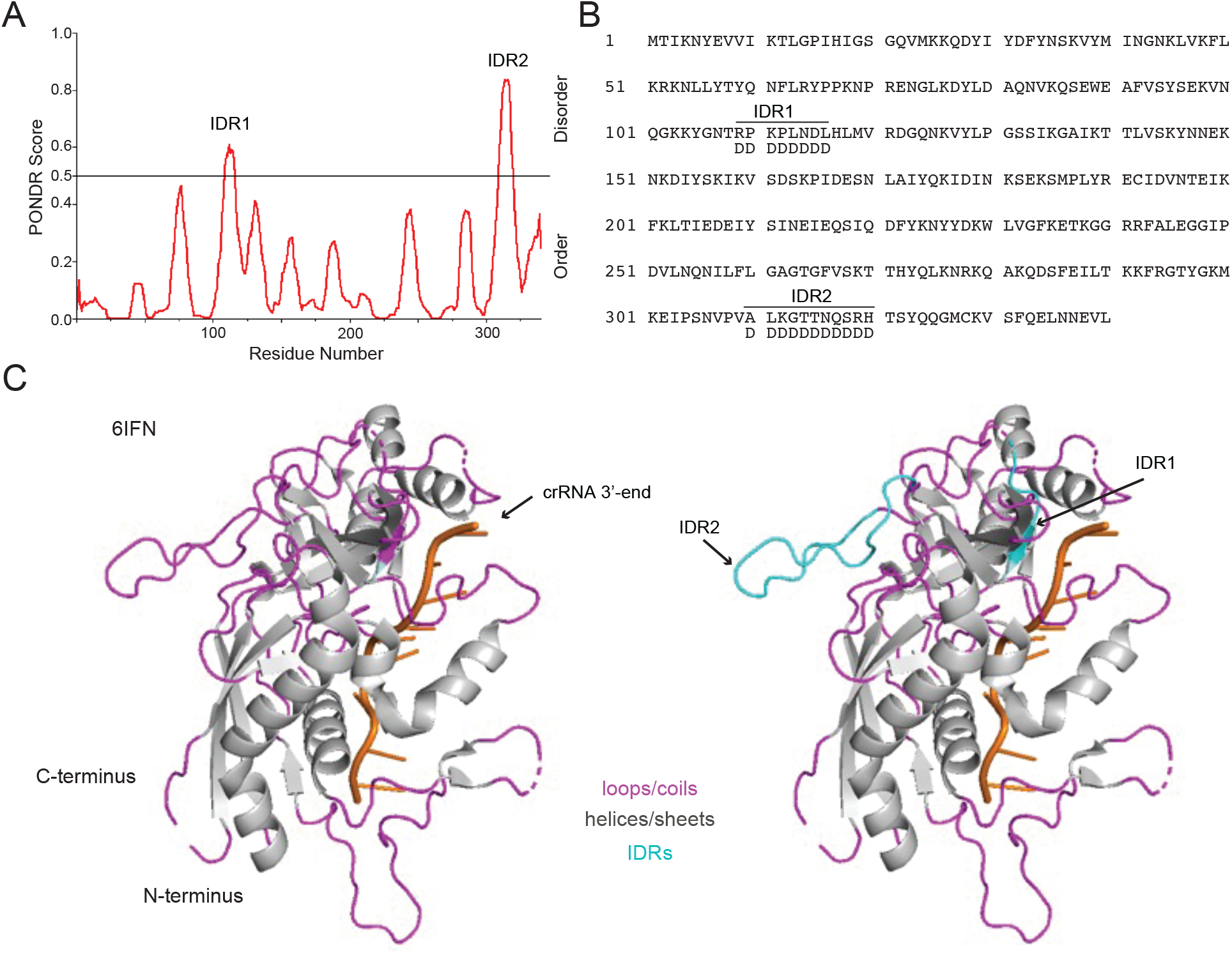
Predicted disordered regions of Csm5. (A) Predicted disordered regions in *S. epidermidis* Csm5 are shown. The PONDR (Predictor of Natural Disordered Regions) score was derived from the web-based tool with the same name (pondr.com). Regions with a score of 0.5 or higher have a high probability of being disordered. The two putative disordered regions of Csm5 are labeled as IDR1 and IDR2. (B) The amino acid sequence of Csm5 is shown with putative disordered regions delimited. IDR1 encompasses residues 109-116, and IDR2 encompasses residues 310-320. (C) The Csm5 subunit of the unbound Cas10-Csm complex from *Streptococcus thermophilus* is shown as determined by (You et al. 2019) (PDB ID 6IFN). Csm5 is colored according to secondary structure with loops/coils in magenta and helices/sheets in grey. The bound crRNA is shown in orange. Homologous residues encompassing predicted IDRs in *S. epidermidis* Csm5 are shown in cyan on the right. The figure was generated using Pymol.

**Figure 5-figure supplement 1.**
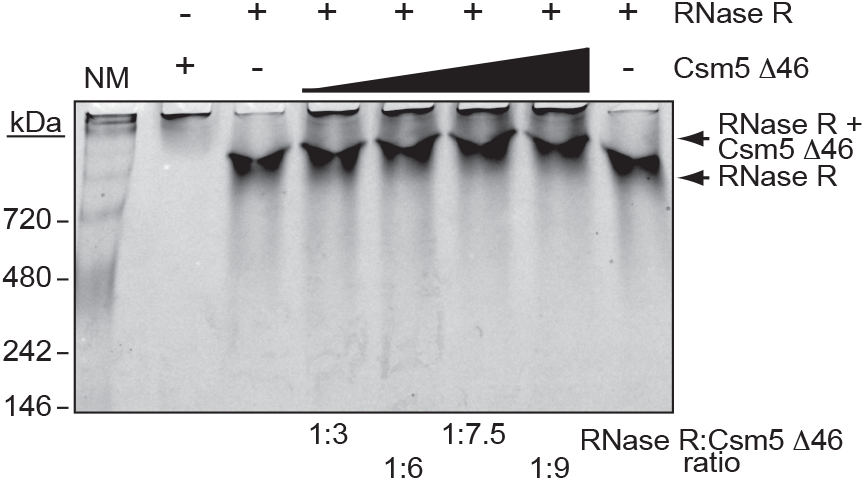
Csm5Δ46 retains interaction with RNase R. Shown is a native gel in which RNase R was resolved alone or with increasing concentrations of Csm5Δ46. The gel shown is a representative of three independent trials. NM, native protein marker; kDa, kilodalton. See Figure 5-figure supplement 1-source data 1.

**Figure 6-figure supplement 1.**
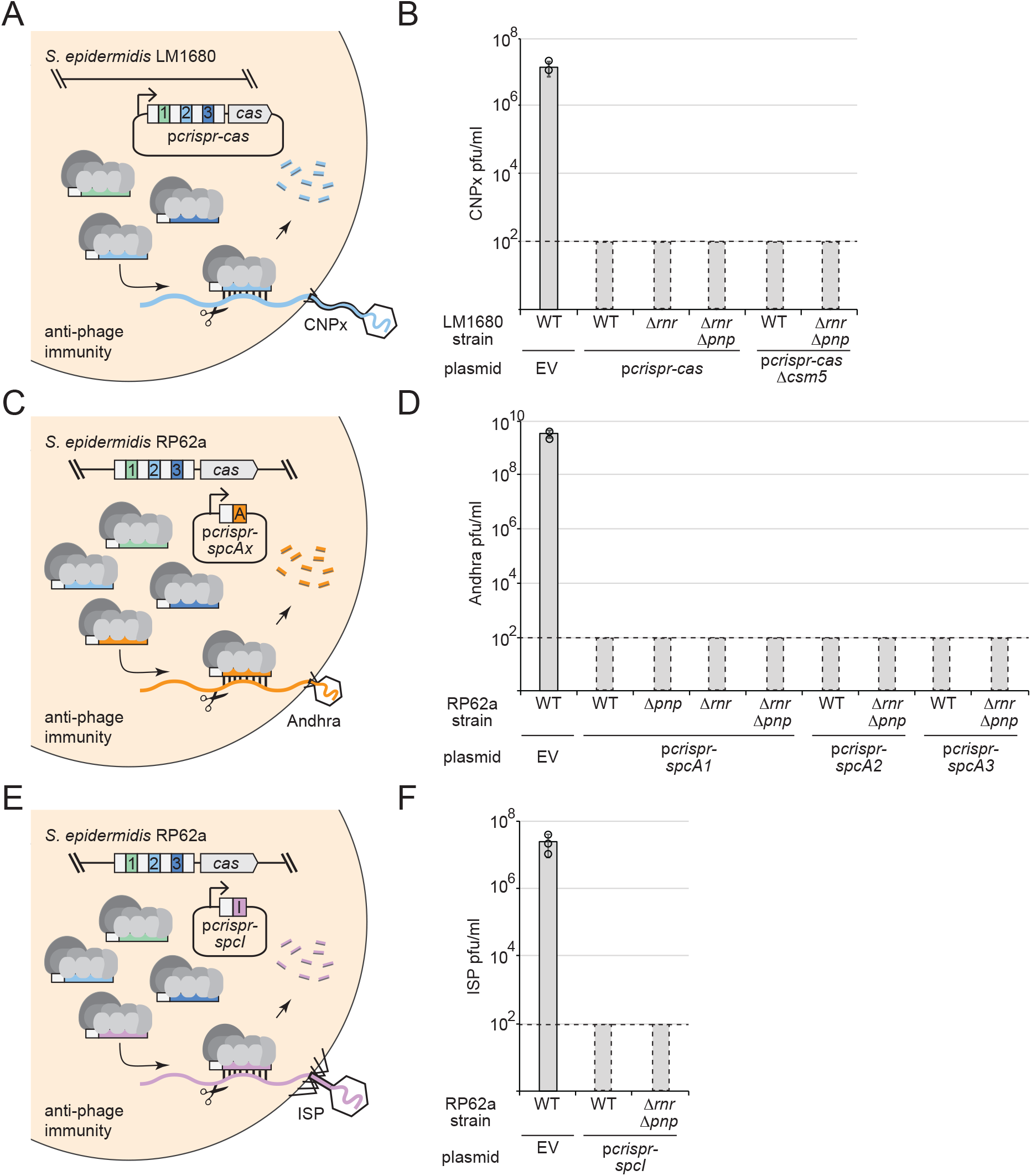
RNase R and PNPase are dispensable for anti-phage immunity. (A) Illustration of the assay used to test anti-phage immunity in *S. epidermidis* LM1680. In this assay, dilutions of the siphophage CNPx are placed atop lawns of cells containing variants of p*crispr-cas*, in which the second spacer (*spc2*, light blue square) bears complementarity to the phage genome. (B) Graph showing results for the anti-phage assay in panel A. Indicated strains were challenged with CNPx and the resulting plaque forming units per milliliter (pfu/ml) were enumerated. See Figure 6-figure supplement 1-source data 1. (C) Illustration of the assay used to test anti-phage immunity in *S. epidermidis* RP62a against phage Andhra. In this assay, the genome-encoded Cas10-Csm complex associates with crRNAs encoded in the plasmid p*crispr-spcAx* (orange square) which target podophage Andhra. (D) Graph showing results for the anti-phage assay in panel C. Each p*crispr-spcA* construct contains a single spacer that targets genes that encode Andhra’s DNA polymerase (*spcA1*), major tail protein (*spcA2*), or lysin-like peptidase (*spcA3*). Indicated strains were challenged with Andhra and the resulting pfu/ml were enumerated. See Figure 6-figure supplement 1-source data 2. (E) Illustration of the assay used to test anti-phage immunity in *S. epidermidis* RP62a against phage ISP. In this assay, the genome-encoded Cas10-Csm complex associates with a spacer encoded in the plasmid p*crispr-spcI* (purple square) which protects against myophage ISP. (F) Graph showing results for the anti-phage assay in panel E. The p*crispr-spcI* construct contains a single spacer that targets ISP’s lysin gene. Indicated strains were challenged with myophage ISP and the resulting pfu/ml were enumerated. See Figure 6-figure supplement 1-source data 3. For all phage challenge assays, the data shown is an average (±S.D.) of three independent trials. Dotted lines on graphs indicate the limits of detection for these assays, and bars with dotted borders were inserted below the line to indicate zero plaques counted. EV, empty vector.

**Table S1.**
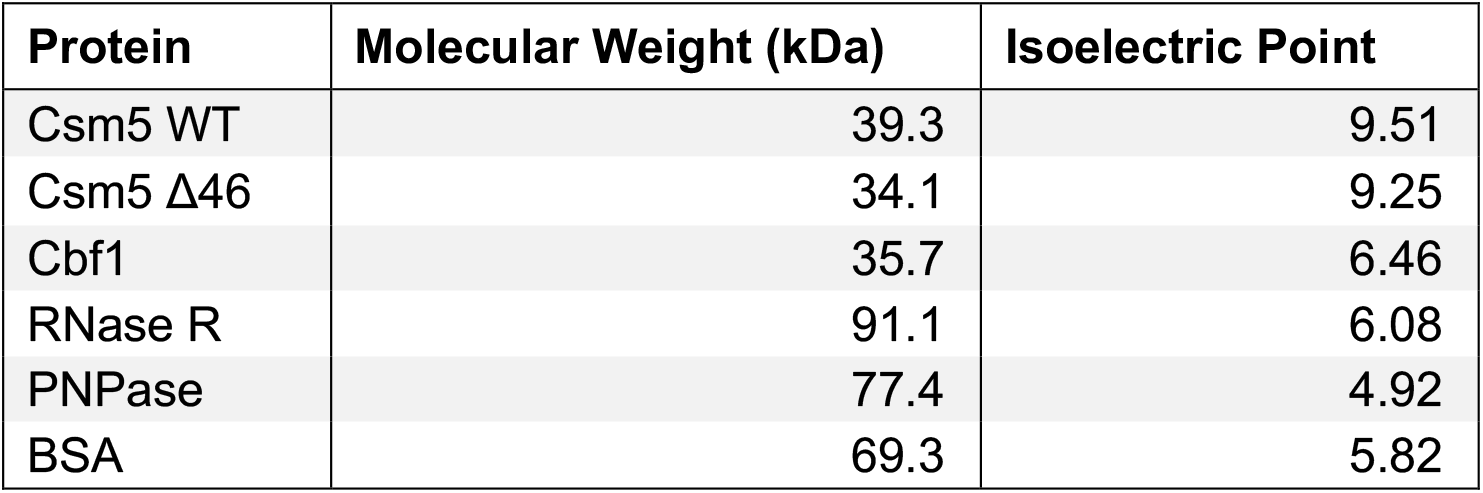
Accompanies Figures 2, 3, 5, Figure 3-figure supplement 1, and Figure 5-figure supplement 1. Theoretical molecular weights and isoelectric points of purified proteins in this study.

**Table S2.**
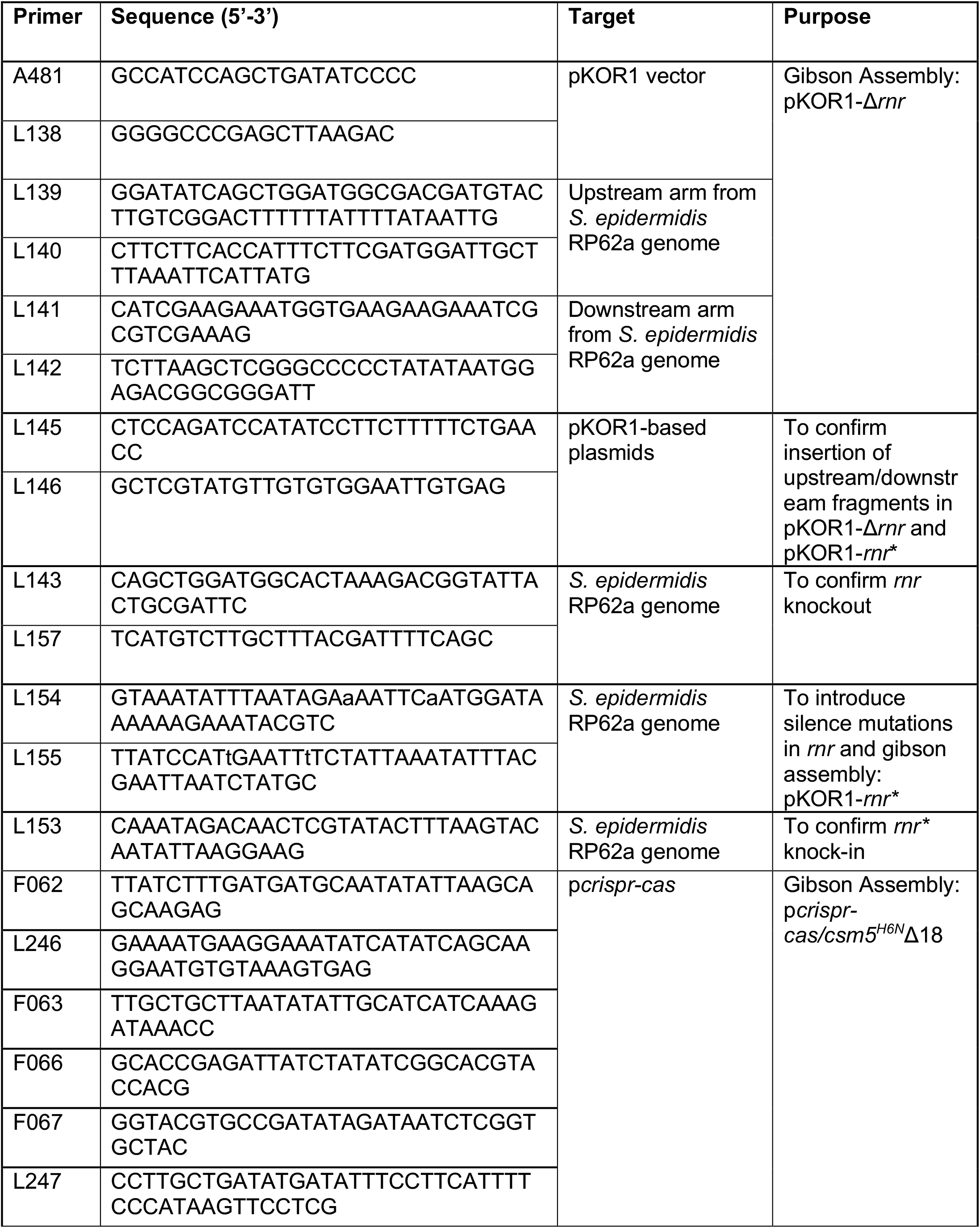
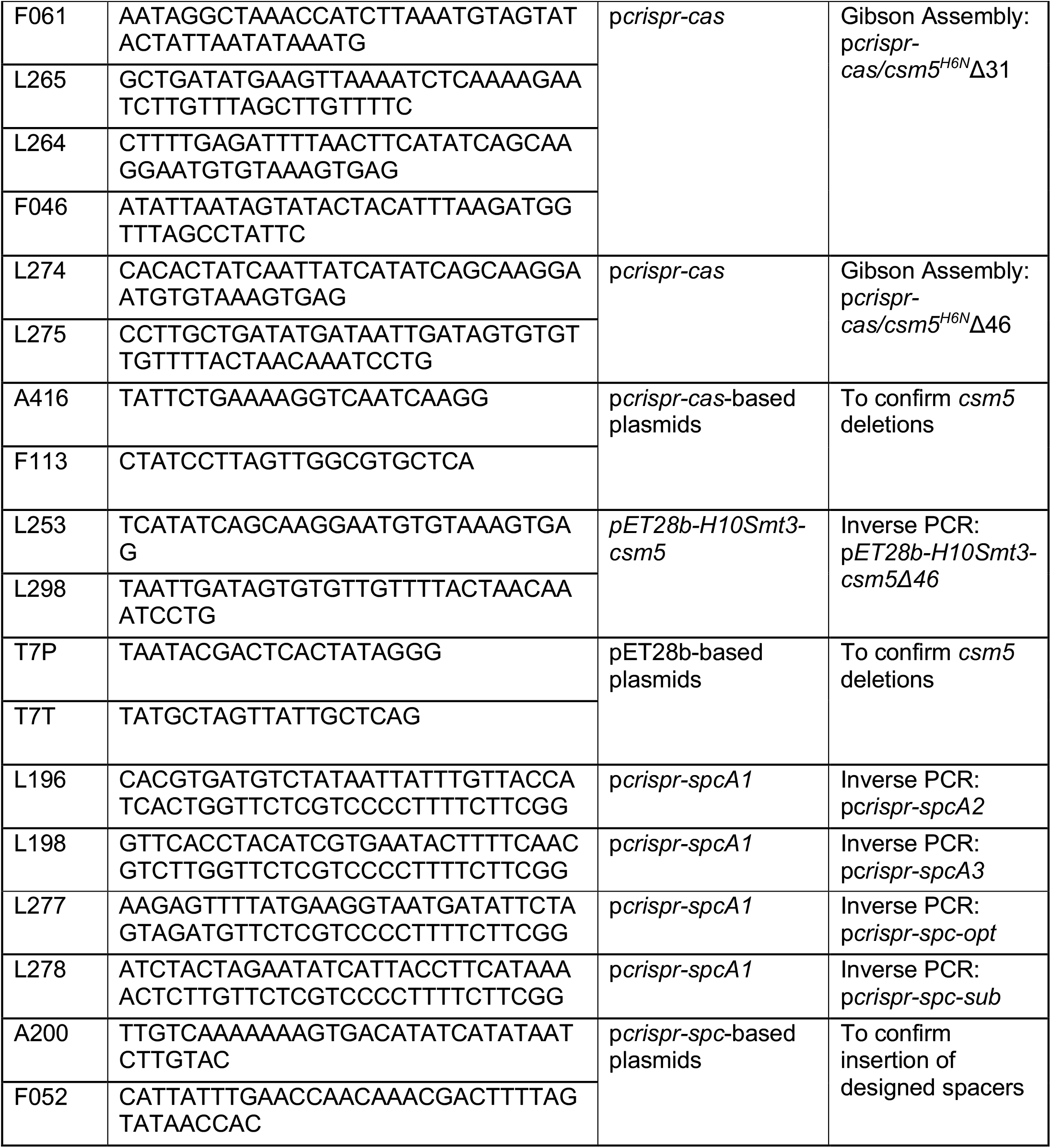
DNA oligonucleotides used for cloning and PCR in this study.

